# Proteome dynamics reveal Leiomodin 1 as a key regulator of myogenic differentiation

**DOI:** 10.1101/2024.03.29.587321

**Authors:** Ellen Späth, Svenja C. Schüler, Ivonne Heinze, Therese Dau, Alberto Minetti, Maleen Hofmann, Katja Hönzke, Julia von Maltzahn, Alessandro Ori

## Abstract

During myogenic differentiation the cellular architecture and proteome of muscle stem cells and myoblasts undergo extensive remodeling. These molecular processes are only partially understood and display alterations in disease conditions as well as during aging resulting in impaired regeneration. Here, we used mass spectrometry to quantify the temporal dynamics of more than 6000 proteins during myogenic differentiation. We identified the actin nucleator leiomodin 1 (LMOD1) among a restricted subset of cytoskeletal proteins increasing in abundance in early phases of myogenic differentiation. We show that LMOD1 is already expressed by muscle stem cells *in vivo* and displays increased abundance during skeletal muscle regeneration, especially during early regeneration suggesting that LMOD1 is important for induction of myotube formation. Of note, knockdown of LMOD1 in primary myoblasts and during skeletal muscle regeneration severely affects myogenic differentiation, while overexpression accelerates and improves the initiation of myotube formation suggesting that LMOD1 is a critical component regulating myogenic differentiation. Mechanistically, we show that LMOD1 physically and functionally interacts with the deacetylase sirtuin1 (SIRT1), a regulator of myogenic differentiation, especially at the onset of myogenic differentiation. We demonstrate that LMOD1 influences SIRT1 localization and the expression of a subset of its target genes. Consistently, depletion or pharmacological inhibition of SIRT1 partially rescues the impairment of myogenic differentiation observed after knockdown of LMOD1. Our work identifies a new regulator of myogenic differentiation that might be targeted to improve muscle regeneration in aging and disease.

## Main Text

Muscle stem cells (MuSCs), also known as satellite cells, possess a remarkable ability to regenerate adult skeletal muscle following injury (Sambasivan et al. 2011; Murphy et al. 2011). These cells are typically quiescent, reside between the basal lamina and the myofiber (Mauro 1961; Sousa-Victor et al. 2015), and are characterized by the expression of the paired box transcription factor 7 (PAX7) (Seale 2003; Lepper et al. 2011; von Maltzahn et al. 2013; Zammit et al. 2006). In response to physiological and regeneration-inducing stimuli, MuSCs become activated, begin to self-renew, and undergo myogenic differentiation (Fu et al. 2015; Schmidt et al. 2019).

During myogenic differentiation after injury - a process recapitulating myogenesis during development - MuSCs proliferate, differentiate, and finally fuse to damaged myofibers or form new myofibers. These processes are tightly regulated by the expression of different myogenic regulatory factors (MRFs) such as MYF5, MYOD, myogenin and MRF4 (Chang and Rudnicki 2014; Schmidt et al. 2019; Tiffin et al. 2003). Additionally, external cues from the surrounding microenvironment and the downstream signalling pathways, including NOTCH, WNT, BMPs, FGFs, and Hedgehog (Fre et al. 2005; Baghdadi, Castel, et al. 2018; Baghdadi, Firmino, et al. 2018; Bentzinger, Wang, von Maltzahn, et al. 2013; Friedrichs et al. 2011; Larraín et al. 1997; Voronova et al. 2013) modulate myogenic differentiation (Bentzinger, Wang, Dumont, et al. 2013; Hung et al. 2023).

Epigenetic mechanisms and chromatin remodeling contribute to this process by influencing the accessibility of transcription factors to regulatory regions of the DNA and, thereby, the expression of myogenic genes (Saccone and Puri 2010; Massenet et al. 2021). Consistently, key epigenetic regulators such as histone deacetylases (HDACs) and methyltransferases, e.g., SIRT1 (Ryall et al. 2015), HDAC4 (Moresi et al. 2012; Finke et al. 2022) and SMYD3 (Codato et al. 2019) have been functionally linked to maintenance of muscle integrity and MuSC differentiation.

Similarly, alterations in mechanosensing and interactions with the extracellular matrix (ECM) were shown to significantly affect the cytoskeletal architecture and function of MuSCs in disease and during aging (Chakkalakal et al. 2012; Hwang and Brack 2018; Lukjanenko et al. 2016). Mutations in cytoskeletal proteins, e.g., dystrophin, plectin or actin binding proteins, impair MuSCs function and are causal to diseases like Duchenne muscular dystrophy (DMD) or Nemaline myopathy (NM) (Winder et al. 1995; Lu-Nguyen et al. 2017; Mournetas et al. 2021; Winter and Wiche 2013; Nguyen et al. 2023). Similarly, age-related changes in cytoskeletal proteins contributing to altered actin dynamics and dysregulated desmin distribution have been implicated in the decline of myofiber functionality and sarcopenia (Meyer et al. 2013; Budai et al. 2018; Larsson et al. 2019; Lai and Wong 2020). Although these mechanisms contribute to remodeling the cellular architecture of MuSCs, a comprehensive understanding of the temporal dynamics of proteome remodeling during differentiation remains lacking. To address this knowledge gap, we performed an unbiased proteomic analysis of the early stages of myogenic differentiation to identify previously unrecognized proteins involved in this process and to examine how they functionally interact with established regulatory pathways. We identified LMOD1 among a restricted group of cytoskeletal proteins that increase in abundance during early myogenic differentiation. LMODs are known as powerful actin filament nucleators but their role in myogenic differentiation has not been investigated so far (Chereau et al. 2008; Boczkowska et al. 2015; Fowler and Dominguez 2017; Tolkatchev et al. 2022). We validated *in vivo* that MuSCs express LMOD1. We showed that LMOD1 is transiently upregulated during skeletal muscle regeneration and that reducing LMOD1 expression after injury impairs skeletal muscle regeneration. Through knockdown and overexpression experiments, we show that LMOD1, but not the closely related LMOD2, is essential for proper myogenic differentiation. Mechanistically, we show that LMOD1 interacts with the deacetylase SIRT1, a known regulator of myogenic differentiation (Fulco et al. 2003; Ryall et al. 2015), and thereby influences its subcellular localization and the expression of a subset of target genes.

### Proteome dynamics during myogenic differentiation

To comprehensively and unbiasedly examine the protein dynamics during myogenic differentiation, we performed an *in vitro* five-day differentiation time course experiment using mouse primary myoblasts from five different mice (Figure 1A) and collected samples at each day of differentiation for label-free quantitative proteomics (Table S1). Principal Component Analysis (PCA) based on protein intensities obtained from mass spectrometry data (6098 quantified proteins in total) revealed a progressive separation of the different cell populations by the day of differentiation (Figure 1B). We confirmed the expected dynamics of key myogenic regulators by qRT-PCR and mass spectrometry (Schmidt et al. 2019; Majchrzak et al. 2024), validating our experimental setup and proper myogenic differentiation (Figures 1C and 1D). Next, we compared protein dynamics by clustering the obtained proteomics data (Table S1). The 3839 protein groups significantly affected in at least one measured time point of differentiation relative to undifferentiated myoblasts were assigned to six clusters by k-means clustering (Figure 1E). Each cluster displayed distinct dynamics and was enriched for specific KEGG pathways (Figure 1F). Notably, cluster 2 included proteins that progressively increased in abundance during differentiation and were closely associated with muscle contraction and function, as well as cytoskeletal dynamics e.g. Actinin Alpha 2 (ACTN2), Tropomyosin alpha-1 and −3 (TPM1, TPM3) Leiomodin 1 (LMOD1) and Dystrophin (DMD) (Figure 1G). These findings confirm the importance of cytoskeletal dynamics during myogenic differentiation but also raise questions about the molecular mechanism responsible for this shift in the proteomic landscape.

**Figure 1:**
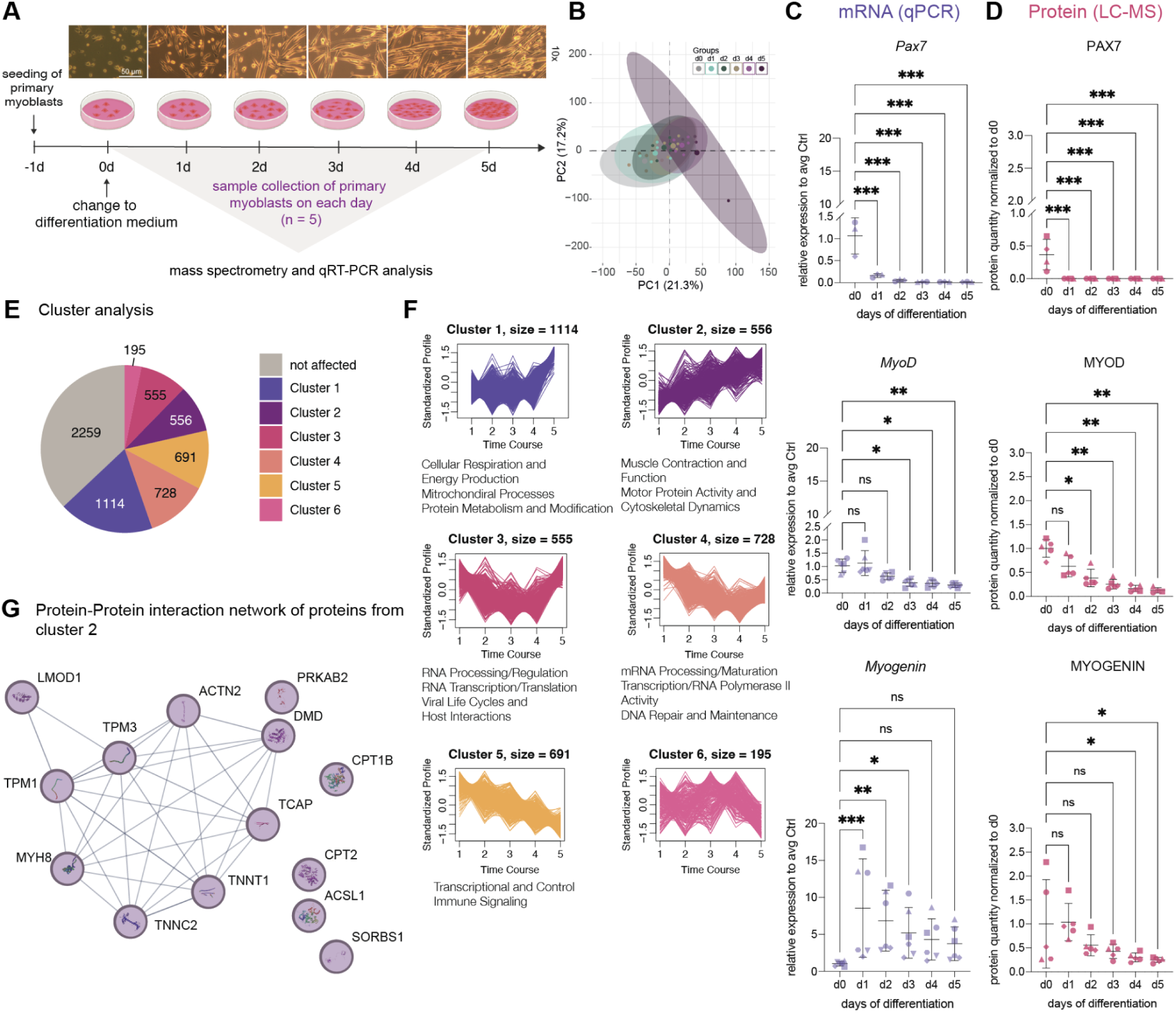
Analysis of the proteome shows dynamic alterations during myogenic differentiation. **A.** Experimental workflow for analyzing the five-day differentiation time course using mass spectrometry (Table S1) and qRT-PCR. Primary myoblasts isolated from five individual mice were seeded and differentiated for up to five days. **B**. Principal component analysis (PCA) of proteomics data. Ellipses represent a 95 % confidence interval for each day of differentiation. **C**. qRT-PCR analysis showing the relative mRNA expression of *Pax7*, *MyoD*, and *Myogenin*, normalized to *Gapdh* and day 0. **D**. Mass spectrometry-based quantification of PAX7, MYOD and MYOGENIN protein levels normalized to day 0. **E**. Pie chart for proteome dynamics. 6098 protein groups were subjected to k-means clustering analysis, 2259 were unaffected, while 3839 protein groups showed significant changes (Absolute AVG log2 ratio>0.58 and Q value<0.25) in expression levels in at least one-time point compared to day 0. Protein groups were classified into six clusters based on their expression dynamics using k-means clustering. **F.** Clusters showing distinct protein abundance dynamics during myogenic differentiation and KEGG pathways enriched (FDR <0.05) in each cluster (Table S1). **G.** Proteins from cluster 2 were chosen according to KEGG pathway annotation: striated muscle contraction and function, motor protein activity and cytoskeletal dynamics. The protein-protein interaction network was visualized with STRING (Szklarczyk et al. 2011), edge confidence: medium 0.4. For (**C**) and (**D**): In all bar plots, each symbol represents a biological replicate, and the error bars indicate the standard deviation. (SD). One-way, ANOVA. *: q value ≤ 0.05, **: q value ≤ 0.01, ***: q value ≤ 0.001.

### LMOD1 levels increase during early myogenic differentiation, in aged MuSCs and regenerating skeletal muscle

Given the role of cytoskeleton remodeling as one of the drivers of myogenic differentiation, we focused on this cluster of proteins and specifically looked for proteins that increased early during the differentiation process. Among these, we found two proteins encoded by paralog genes, namely leiomodin 1 (LMOD1) and leiomodin 2 (LMOD2) (Figure 2A), which share 67.05 % sequence similarity, although varying in their number of amino acids and expression across tissues (Qualmann and Kessels 2009; Fowler and Dominguez 2017). Interestingly, the expression pattern of LMOD1 and LMDO2 during myogenic differentiation displayed differences: while LMOD1 showed a significant increase at the beginning (day 1) (Figure S1C) of myogenic differentiation, LMOD2 increased later in the process (Figure 2A), suggesting that LMOD1 might be a key protein in driving early myogenic differentiation. Surprisingly, we detected LMOD1 in freshly isolated muscle stem cells (MuSCs), but not LMOD2. Additionally, we observed that the protein levels of LMOD1 increased in MuSCs isolated from older mice (Figure 2C and Figure S1B). We further analyzed published transcriptomic data sets that describe changes between young and old MuSCs in both quiescent and activated states in young and old animals (Liu et al. 2013; Lukjanenko et al. 2016). In these analyzed transcriptomic data sets, Lmod1 was found to be significantly downregulated during the activation of MuSCs in both young and old mice (see Figure S1B). To assess the *in vivo* relevance of our finding, we queried two proteomic datasets of freshly isolated MuSCs and four different skeletal muscles (gastrocnemius, G; soleus, S; tibialis anterior, TA; extensor digitorum longus, EDL) (Schüler et al. 2021). We found LMOD2 to be the most abundant leiomodin protein in whole skeletal muscle, consistent with data from (Tsukada et al. 2010; Nworu et al. 2015; Kiss et al. 2020), while the overall abundance of LMOD1 was lower since this protein has been mainly associated with smooth muscle cells (Nanda and Miano 2012; Conley et al. 2001; Nanda et al. 2018) (Figure 2B). To independently validate this finding, we performed immunofluorescence stainings on cryosections of TA muscles from young and geriatric mice. Consistent with previous publications, we observed a reduction in the number of PAX7+ cells with increasing age (Brack et al. 2005; Shefer et al. 2006; Yamakawa et al. 2020) (Figure S1C). Furthermore, we confirmed the age-related increase of LMOD1 abundance in MuSCs (Figure 2D and Figure S1B). Of note, this was accompanied by a significant increase in the number of LMOD1+/PAX7+ cells in sections of geriatric compared to young mice (Figure 2E). Interestingly, we found LMOD1 to be mainly located in the cytoplasm of PAX7+ cells in TA muscles from young mice (Figure 2D and S1D). However, in TA muscles from geriatric mice, we found a significantly higher number of PAX7+ cells that displayed nuclear localization of LMOD1 (Figure 2F), suggesting that LMOD1 accumulates and mislocalizes in MuSCs with increasing age and could be one contributing factor to the reduced ability of MuSCs to differentiate in aged mice.

**Figure 2:**
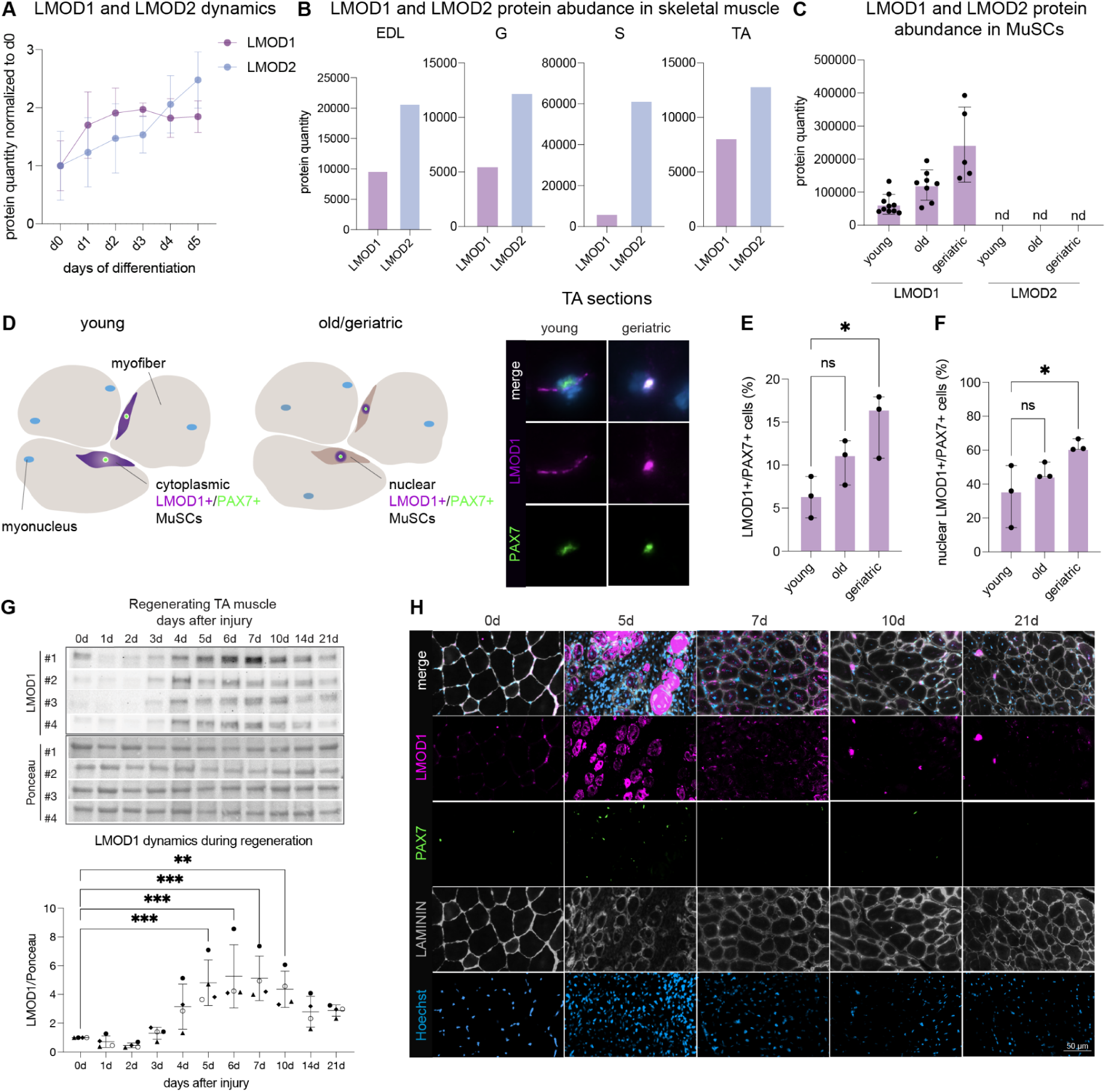
LMOD1 increases during early myogenic differentiation and is upregulated in aged MuSCs. **A.** Protein quantification of LMOD1 and LMOD2 during myogenic differentiation, based on mass spectrometry data and normalized to the protein amount of day 0. Error bars indicate SD, and colored dots indicate the mean value of the biological replicates (n = 5). **B.** LMOD1 and LMOD2 protein abundance estimated from mass spectrometry data in different skeletal muscles (gastrocnemius, G; soleus, S; tibialis anterior, TA; extensor digitorum longus, EDL). Displayed values are averages of n = 5 samples from individual mice. Mass spectrometry data from (Schüler et al. 2021). **C**. LMOD1 and LMOD2 protein quantity in MuSCs obtained from young (n = 10), old (n = 8) and geriatric (n = 5) mice, nd = not detected. Mass spectrometry data from (Schüler et al. 2021). **D**. Representative immunofluorescence images of LMOD1 (purple), PAX7 (green), and Hoechst (blue) of TA sections from n = 3 individual mice per age group: young (3-month-old) and old (18-months-old), geriatric (33-months-old) mice. Scale bar: 5 µm. **E.** Ratio of PAX7+ and LMOD1+ cells normalized to the cells positive for PAX7+ (Figure S2C) and **F.** Quantification of the percentage of PAX7+ cells showing nuclear LMOD1 localization. Related to Figure S1. For (**E**) and (**F**): One-way, ANOVA. *: q value ≤ 0.05, **: q value ≤ 0.01, ***: q value ≤ 0.001. In all bar plots, each black dot represents a biological replicate, and the error bars indicate the SD. **G**. Muscle regeneration time course: Immunoblot of LMOD1 levels in regenerating TA muscle after injury. Quantification of LMOD1 levels were normalized to the respective entire lane of the Ponceau staining and relative to the LMOD1 signal at day 0. Each symbol represents a biological replicate, and error bars represent SD. One-way ANOVA; **: p-value ≤ 0.01, ***: p-value ≤ 0.001. **H.** Representative immunofluorescence images of LMOD1 (purple), PAX7 (green), LAMININ (grey), and Hoechst (blue) of TA sections at different days post-injury from young mice (3 months old). Scale bar: 50 µm.

Finally, we investigated the expression of LMOD1 during regeneration of skeletal muscle *in vivo.* Therefore, we performed a regeneration time course experiment following cardiotoxin (CTX) injury and assessed LMOD1 expression by immunoblot and immunofluorescence analyses in young mice (Figure 2G and Figure 2H). We observed low abundance of LMOD1 in uninjured TA muscle, but found a progressive increase in abundance up to day 7 after injury and a decrease thereafter (Figure 2G). Immunofluorescence staining confirmed that LMOD1 was highly abundant in myofibers at day 5 after injury, when mostly newly formed regenerating myofibers are present, hinting at a functional role of LMOD1 in early myogenic differentiation. Of note, LMOD1 abundance was barely detectable at day 7 and later time points, suggesting that LMOD1 abundance decreases during maturation of myofibers (Figure 2H). In conclusion, these findings demonstrate that LMOD1 abundance is not only increased at the onset of myogenic differentiation *in vitro* but is also expressed by MuSCs under homeostatic conditions *in vivo* and in MuSC and newly formed myofibers following injury, indicating its importance for myofiber differentiation and skeletal muscle regeneration.

**Figure S1:**
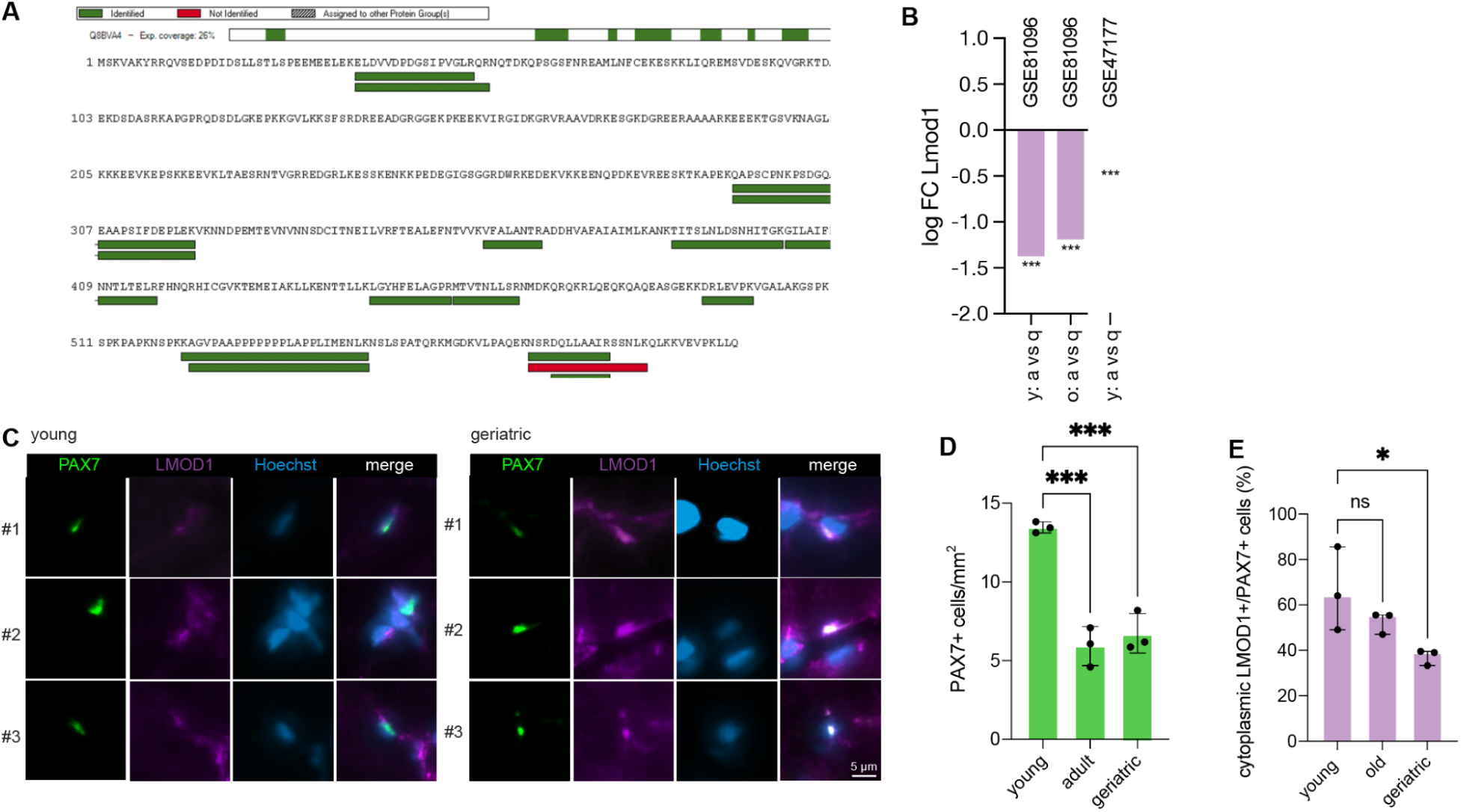
LMOD1 in MuSCs and skeletal muscle. **A.** LMOD1 protein coverage in proteomics data from MuSCs. Multiple unique (proteotypic) peptides were identified. Data from (Schüler et al. 2021). **B.** Barplot showing log2 fold changes of Lmod1 mRNA abundance in activated vs. quiescent MuSCs (in young and old) from previously published transcriptome data sets (GSE81096) (Lukjanenko et al. 2016) and GSE47177 (Liu et al. 2013). *: adjusted p-value < 0.05, ***: adjusted p-value < 0.001; a - activated, q - quiescent, o - old, y - young. C. Immunofluorescence images of TA sections from different mice (n = 3) per age group of young (3 months old) and geriatric (33 months old) mice. PAX7 (green), LMOD1 (purple) and Hoechst (blue). **D**. Number of PAX7+ cells per mm^2^ cells in muscle sections from young, old and geriatric mice and **E**. Quantification of PAX7+ cells showing cytoplasmic LMOD1 localization. For (**D**) and (**E**): One-way ANOVA; *: p-value ≤ 0.05, ***: p-value ≤ 0.001, ns: not significant. In all bar plots, each black dot represents a biological replicate, and the error bars indicate the SD. Related to Figure 2.

### Knockdown of *Lmod1* impairs myogenic differentiation, while overexpression of *Lmod1* accelerates myotube formation

The early increase of abundance of LMOD1 during myogenic differentiation suggests a potential functional role for this protein in promoting the initiation of myogenic differentiation. To test this hypothesis, we used a small interfering RNA (siRNA) to reduce LMOD1 expression and analyzed the impact on myoblast proliferation (Figure S2A) and differentiation (Figure 3A). We confirmed the successful knockdown of *Lmod1* mRNA by qRT-PCR (Figure S2B and S2F) and reduced levels of LMOD1 protein by immunoblot analysis (Figure S2C and S2G). After 48 hours of proliferation, we observed a significant 26% decrease in the number of proliferating myoblasts (Ki67+/Hoechst+) upon knockdown of LMOD1 (Figure S2D and S2E). This was accompanied by a proportional increase in non-proliferating myoblasts (Ki67-/Hoechst+), while the total Hoechst-positive cell count (Ki67+/Hoechst+ and Ki67-/Hoechst+) remained unchanged (Figure S2E).

**Figure 3:**
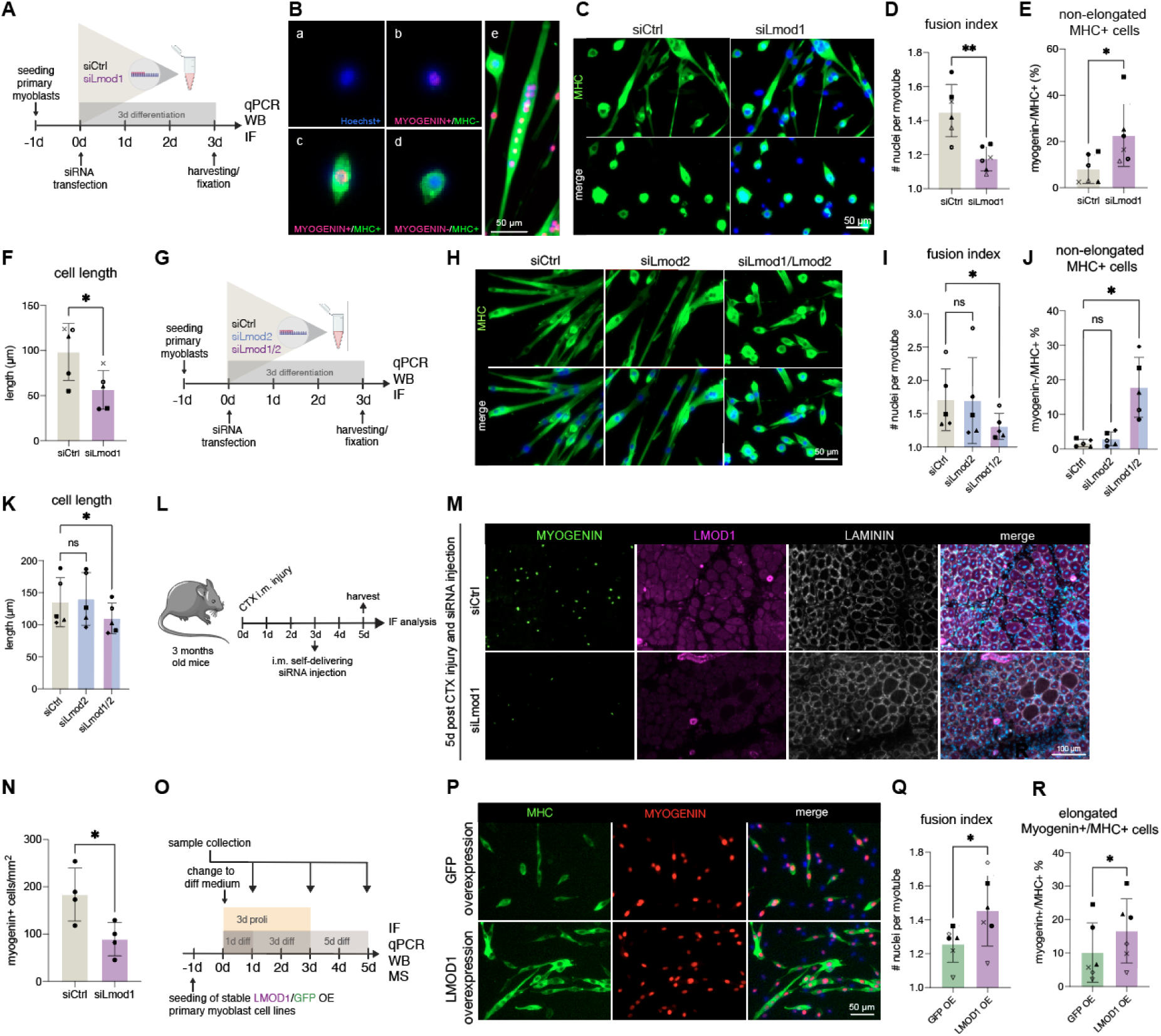
Knockdown of *Lmod1* impairs myogenic differentiation and reduces muscle regeneration, while overexpression enhances it. **A.** Schematic of the *Lmod1* knockdown experiment after 3 days of differentiation. siRNA directed against *Lmod1* or scramble (siCtrl) control was used to transfect primary myoblasts isolated from individual mice. Differentiation was induced by a change to differentiation medium at 0h. **B.** Overview of quantified different cell types based on their expression of myogenic markers; Myogenin (red), Myosin heavy chain (MHC) (green), Hoechst (blue). Scale bar: 50 µm. (a) non-proliferating myoblasts: Hoechst+/Myogenin-/MHC- and undifferentiated. (b) Myogenin+/MHC-: Myogenin positive, MHC negative, just started to differentiate. (c) Myogenin+/MHC+: co-express bot in the process of differentiation. (d) Myogenin-/MHC+: non-elongated. (e) Fully differentiated myotubes: quantified by nuclei count to assess the fusion process. **C.** Representative immunofluorescence images after three days of differentiation and siCtrl or siLmod1 transfection; MHC (green), nuclei (blue). Scale bar: 50 µm. **D - F.** Quantification of the number of nuclei per myotube (**D**), Myogenin-/MHC+ cells (**E**) and length of differentiated myotubes (in µm) (**F**). Paired t-test; *: p-value ≤ 0.05, **: p-value ≤ 0.01. **G.** Schematic of the *Lmod2* and *Lmod1/Lmod2* double knockdown experiment after 3 days of differentiation. siRNA against *Lmod1*, *Lmod2*, both or scramble (siCtrl) was used to transfect primary myoblasts. Differentiation was induced by a change to differentiation medium at 0h. **H.** Representative immunofluorescence images of siLmod2 and siLmod1/Lmod2 double knockdown after three days of differentiation; devMHC (green), nuclei (blue). Scale bar: 50 µm. **I - K.** Quantification of the number of nuclei per myotube (**I**), Myogenin-/MHC+ cells (**J**), Length of differentiated myotubes (in µm) (**K**). One-way ANOVA; *: p-value ≤ 0.05, ns: not significant. **L**. Experimental schematic for analysis of *in vivo* CTX-induced injury of TA muscles combined with injection of self-delivering siRNAs at 3 days post injury. n = 4 mice per group, 3 months old. **M**. Representative immunofluorescence images of MYOGENIN (green), LAMININ (grey), and Hoechst (blue) of TA-sections 5 days post-injury from young mice (3 months old). **N.** Quantification of the number of Myogenin+ cells normalized per area. Unpaired t-test; *: p-value ≤ 0.05. **O.** Illustration of the experimental setup. Primary myoblasts isolated from individual mice stably overexpressing (OE) LMOD1 (purple) or GFP (green) were seeded and either collected during proliferation (3d proli), after 1 day (1d diff), 3 days (3d diff) or 5 days (5d diff) of differentiation. Differentiation was induced by a change to differentiation medium at 0h. Cells were harvested for immunofluorescence analysis (IF), qRT-PCR, immunoblot (WB) or mass spectrometry (MS). **P.** Representative immunofluorescence images of primary myoblasts after one day of differentiation, showing either stable expression of LMOD1 or GFP. devMHC (green), MYOGENIN (red) and nuclei (blue). Scale bar is 50 µm. **Q** and **R.** Quantification of nuclei per myotube of GFP OE or LMOD1 OE cells (**Q**) and cells expressing Myogenin+/MHC+ defined as just differentiated cells (**R**) after 1 day of differentiation. Paired t-test *: p-value ≤ 0.05. Related to Figure S2. In all bar plots, each symbol represents a biological replicate, and the error bars indicate the SD.

During myogenic differentiation, the cell population becomes heterogeneous with a distinct expression of myogenic markers (such as MyoD, myogenin, and devMHC) determining cellular identity thereby controlling the potential to differentiate and ultimately form new myofibers (Figure 3B). We characterized the cell populations after knockdown of LMOD1 based on the expression of myogenic markers using immunofluorescence staining at day three of differentiation (Figure 3C and S2H). We found that the reduced levels of LMOD1 resulted in an increased frequency of cells with a pronounced spherical phenotype, also identified as non-elongated, MHC-positive cells (Figure 3C and 3E). Moreover, a significant reduction in the number of nuclei per myotube was observed upon knockdown of LMOD1, indicating a potential role in the fusion process (Figure 3D). Furthermore, we observed significantly shorter myotubes upon knockdown of LMOD1, supporting our previous finding that the cells are unable to properly differentiate (Figure 3F).

To determine whether the observed effects were due to a specific functional disruption rather than a decline in cell viability, we conducted a TUNEL assay (Figure S2I). The results revealed that there was no increase in apoptosis after knockdown of *Lmod1*, suggesting that LMOD1 plays an active role in the structural reorganization necessary for myogenesis, rather than being critical for cell survival.

Given the structural similarities between LMOD1 and LMOD2, we set out to determine if individual knockdown in primary myoblasts induces similar phenotypes during myogenic differentiation (Figure 3G). Knockdown of both *Lmod1* and *Lmod2* mRNA was validated by qRT-PCR (Figure S2J) and confirmed at the protein level by immunoblot (Figure S2K). Interestingly, the knockdown of LMOD2 did not lead to any noticeable phenotypes compared to primary myoblasts transfected with a scramble control siRNA (Figure 3H and S2L). However, we observed a significant reduction in the number of nuclei per myotube (Figure 3I), increased frequency of non-elongated MHC-positive cells (Figure 3J), and increased number of shorter myotubes (Figure 3K) in the *Lmod1/2* double knockdown. These data suggest that LMOD1 is required for myogenic differentiation in primary myoblasts, while LMOD2 appears to be dispensable.

To examine how reduced levels of LMOD1 affect skeletal muscle regeneration *in vivo*, we injured the TA-muscle of 3-months-old mice using CTX, followed by injection of a self-delivering siRNA targeting Lmod1 (Figure 3L) at day 3 after injury, a time point when MuSCs start to differentiate. Subsequently, TA-muscles were isolated at day 5 after injury, the time when LMOD1 was previously shown to be highly expressed in newly formed myofibers. Knockdown of *Lmod1* led to a significant reduction in the number of cells expressing the early differentiation marker myogenin (Figure 3M and 3N), suggesting a critical role for LMOD1 in the transition of MuSCs from proliferation to differentiation. Together, these findings indicate that LMOD1 is not only important for myofiber formation but may also play a key regulatory role in the differentiation of MuSCs into myoblasts and myocytes during the muscle regeneration process, stressing its role in controlling myogenic differentiation.

To test whether LMOD1 alone is sufficient to promote myogenic differentiation, we generated six independent primary myoblast cultures that were stably transduced with a virus to overexpress either LMOD1 or GFP (Figure 3O). Overexpression (OE) of Lmod1 mRNA by qRT-PCR (Figure S2M) and protein by immunoblot (Figure S2N) was confirmed by comparison to a GFP OE control. Immunofluorescence analysis (Figure 3P and S2P) after one day of differentiation showed that overexpression of LMOD1 promotes myogenic differentiation, as shown by a higher number of nuclei per myotube (Figure 3P). Additionally, we observed an accelerated formation of fully differentiated myotubes (Figure 3Q) and an increase in myotube length (Figure S2O) in LMOD1 OE cells compared to the GFP OE control construct. To further characterize the molecular alterations induced by overexpression of LMOD1 during myogenic differentiation, we generated proteomics data from LMOD1 overexpressing and GFP control cells at multiple time points of differentiation (Table S2). Using gene set enrichment analysis, we found that at day 1 of differentiation, cells overexpressing LMOD1 exhibit higher levels of proteins that are up-regulated at the beginning of differentiation, such as HSPB3, ALDH3A1, SMTNL2, and ACTG1, when compared to the GFP control group (Figure S2Q) (Table S2). This proteome signature corroborates the observation that LMOD1 overexpression is sufficient to promote myogenic differentiation.

**Figure S2:**
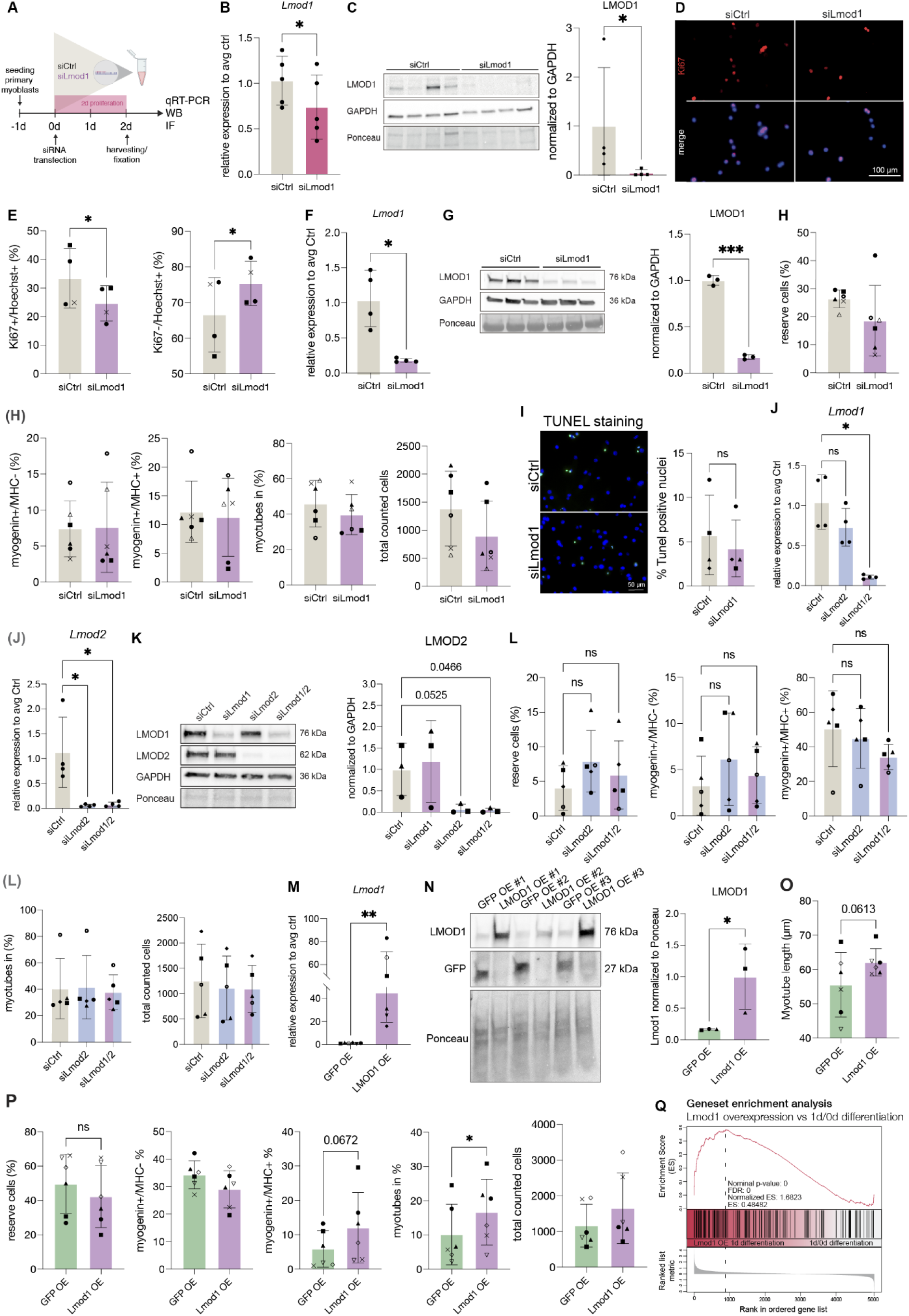
Validation of knockdown efficiency and statistical analysis of siLmod1/Lmod2 knockdown and LMOD1 overexpression. **A.** Schematic of the *Lmod1* knockdown experiment under proliferating conditions. siRNA directed against *Lmod1* or scramble (siCtrl) control was used to transfect primary myoblasts isolated from individual mice. **B.** qRT-PCR analysis showing the relative expression of *Lmod1* after siLmod1 knockdown compared to siCtrl transfected cells, normalized to *Gapdh* expression levels. **C.** Immunoblot and quantification of LMOD1 after transfections with siLmod1 and siCtrl and two days of proliferation normalized to GAPDH. Paired t-test: *: p-value ≤ 0.05. **D.** Immunofluorescence analysis of primary myoblasts stained for the proliferation marker Ki67 (red) and Hoechst (blue). Scale bar: 100 µm. **E** Quantification of cell populations identified by immunofluorescence staining: non-proliferating myoblasts being Hoechst+/Ki67- and cells being Hoechst+/Ki67+ in siCtrl and siLmod1 transfected conditions and the total counted cells per condition. **F.** and **G.** Validation of the siLmod1 knockdown compared to siCtrl transfected cells under differentiation conditions. qRT-PCR for *Lmod1* normalized to *Gapdh* expression levels (**F**) and immunoblot analysis with quantification of LMOD1 signal normalized to GAPDH (**G**). Paired t-test *: p-value ≤ 0.05, ***. **H.** Quantification of cell populations identified by immunofluorescence staining: non-proliferating myoblasts/reserve cells (Hoechst+/Myogenin-/MHC-) cells, Myogenin+/MHC-cells, Myogenin+/MHC+, fully differentiated myotubes and total counted cells under differentiating conditions. **I.** Representative images of primary myoblasts transfected with a non-targeting control siRNA (siCtrl) or an siRNA targeting Lmod1 (siLmod1). Apoptotic cells were identified by TUNEL staining (green), and all nuclei were counterstained with Hoechst (blue). Bar graph showing the percentage of TUNEL-positive nuclei. **J.** and **K.** Validation of the *siLmod1* or *siLmod2* (single) or *siLmod1/Lmod2* (double) knockdown compared to siCtrl transfected cells under differentiation conditions. qRT-PCR for *Lmod1* and *Lmod2* normalized to *Gapdh* expression levels (**J**) and representative immunoblot with quantification of LMOD2 normalized to GAPDH (**K**). One-way ANOVA *: p-value ≤ 0.05 or numbers are indicated. ns: not significant. **L.** Quantification of cell populations (in %) identified by immunofluorescence staining after siLmod1 (single), siLmod2 (single) and siLmod1/2 (double) knockdown under differentiating conditions: non-proliferating myoblasts (Hoechst+/Myogenin-/MHC-) cells, Myogenin+/MHC-cells, Myogenin+/MHC+, fully differentiated myotubes and total counted cells under differentiating conditions. Related to Figures 3I and 3J. **M** and **N.** Validation of LMOD1 OE compared to GFP OE cells after one day of differentiation. Relative *Lmod1* expression assessed by qRT-PCR was compared to GFP OE cells and normalized to *Gapdh* expression levels (**M**) and immunoblot with quantification of LMOD1 normalized to GAPDH (**N**). Paired t-test: *: p-value ≤ 0.05. **O** and **P.** Quantification of cell populations identified by immunofluorescence staining after LMOD1 OE compared to GFP OE: non-proliferating myoblasts (Hoechst+/Myogenin-/MHC-) cells, Myogenin+/MHC-cells, Myogenin+/MHC+, fully differentiated myotubes, total counted cells after and myotube length (in µm) (**O**). Paired t-test: numbers are indicated, ns: not significant. Related to Figures 3N and 3O. **Q.** Gene set enrichment analysis (GSEA) was based on a gene set containing proteins that significantly increased at one day of differentiation compared to 0h/proliferating primary myoblasts (AVG Log2 Ratio > 0.58 and Q value < 0.05, 189 proteins) from the proteomic data generated in (Figure 1A). The GSEA was performed using this gene set on the comparison LMOD1 OE vs. GFP OE at 1 day of differentiation (Table S2). In all bar plots, each symbol or black dot represents a biological replicate, and the error bars indicate the SD.

### LMOD1 interacts with SIRT1 and SIRT2

To elucidate the mechanism by which LMOD1 contributes to myogenic differentiation, we performed a proximity-dependent biotinylation experiment (BioID) for LMOD1 to identify its interaction partners in HEK293T cells (Figure 4A). Given their sequence similarity but distinct functions during myogenic differentiation, we analyzed the interaction partners of LMOD1 and LMOD2 in parallel and focused on hits retrieved exclusively by LMOD1. Therefore, we fused the promiscuous biotin ligase (BirA*) C-terminal to Lmod1 or Lmod2. A cell line expressing only the sequence for BirA* was used as a control (BirA*-Ctrl) to account for non-specific biotinylation. The tetracycline-dependent expression of the respective fusion proteins was validated by immunofluorescence (Figure 4B and 4C) and anti-FLAG immunoblot (Figure S3A). Biotinylation activity, after the addition of exogenous biotin, was assessed using streptavidin-HRP immunoblot (Figure S3A) and immunofluorescence staining (Figure 4B and 4C). Subsequently, biotinylated proteins were captured using streptavidin enrichment and analyzed by mass spectrometry (Table S3). PCA analysis demonstrated a distinct separation between C-terminal fused LMOD1 and LMOD2 expression constructs (Figure 4D and Figure S3D). Using both LMOD1 and LMOD2, we found a significant enrichment compared to the BirA*-Ctrl for known interaction partners (Fowler and Dominguez 2017; Boczkowska et al. 2015) involved in actin filament nucleation such as Tropomodulins (TMODs) and Tropomyosins (TPMs) (Figure S3B-C and Figure S3E), validating our constructs. Interestingly, a direct comparison of streptavidin-enriched proteins from LMOD1-BirA* vs. LMOD2-BirA* revealed a subset of interaction partners that specifically interact with LMOD1. These include the histone deacetylases sirtuin 1 (SIRT1) and sirtuin 2 (SIRT2), as well as the RuvB Like AAA ATPase 1/2 (RUVBL1/2) (Figure 4E and 4F). Sirtuins are involved in various biological processes, including DNA regulation, metabolism, longevity, and myogenesis (Horio et al. 2011; Abdel Khalek et al. 2014; Vachharajani et al. 2016). In particular, one study revealed that mice with muscle-specific inactivation of Sirt1 displayed repression of the myogenic program (Ryall et al. 2015). Since SIRT1 was already identified as a potential mediator of myogenesis, we wanted to validate its preferential interaction with LMOD1 by co-immunoprecipitation independently. Therefore, we transfected SIRT1 as a GFP fusion protein in HEK293T expressing LMOD1-BirA* or LMOD2-BirA* and performed pull-down experiments using a GFP trap. Immunoblot analysis of the GFP-trap eluates using anti-BirA antibodies confirmed a co-precipitation between LMOD1 and SIRT1, while LMOD2 co-precipitated with SIRT1 to a lower extent (Figure 4G). To further validate the interaction of LMOD1 and SIRT1 in primary myoblasts and myogenic differentiation, we performed *in situ* detection of LMOD1 and SIRT1 using proximity ligation assays (PLA) (Figure 4H). In proliferating myoblasts, when the abundance of LMOD1 is low, only a weak PLA signal indicating close proximity of SIRT1 and LMOD1 was detectable both in the cytoplasm and the nucleus. However, at the beginning of myogenic differentiation, when the abundance of LMOD1 increases, an increase in PLA signals indicating close proximity of LMOD1 and SIRT1 was detectable almost exclusively in the cytoplasm. Together these data show that LMOD1 preferentially interacts with the deacetylase SIRT1, suggesting a potential functional interaction between these two proteins in modulating myogenic differentiation. This potential functional role is further supported by our *in vivo* findings in regenerating muscle, where we observed a parallel increase in both LMOD1 and SIRT1 abundance by day 5 post-injury (Figure 4I), a time point critical in the formation of new myofibers.

**Figure 4:**
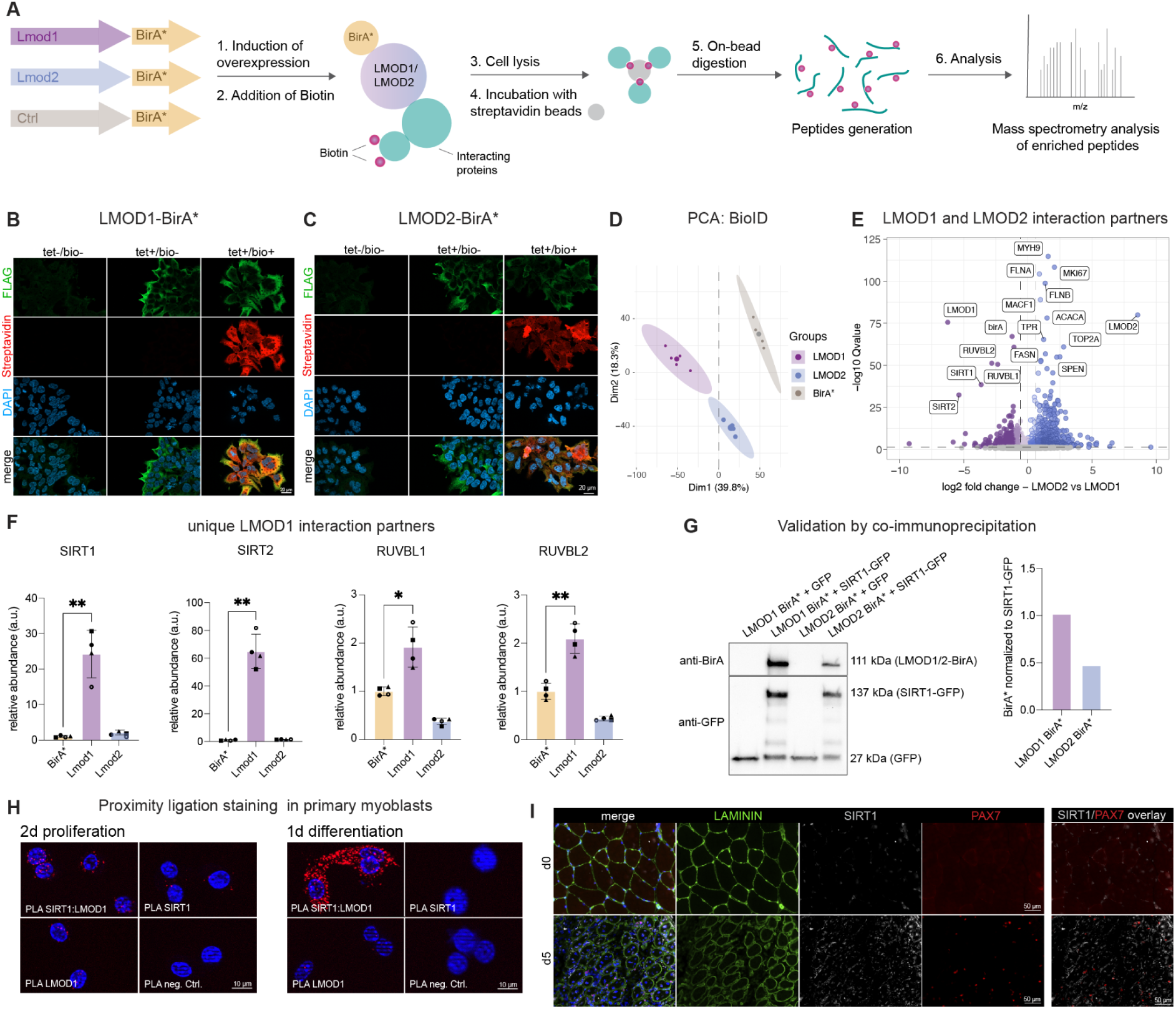
LMOD1 interacts with SIRT1. **A.** BioID workflow. Lmod1 and Lmod2 were C-terminally fused to a promiscuous biotin ligase (BirA*) and expressed in HEK293T cells. BirA* alone served as a control (Ctrl) to assess non-specific biotinylation. Overexpression of fusion proteins was induced by addition of tetracycline. Exogenous biotin was introduced to label interaction partners in close proximity. Biotinylated proteins were captured using streptavidin enrichment, followed by mass spectrometry analysis to identify and quantify proximal interactors. **B.** and **C.** Immunofluorescence analyses of LMOD1-BirA*-FLAG and LMOD2-BirA*-FLAG in HEK293T cells 4 days after seeding without addition of any substance (-tet/-bio), with addition of only tetracycline for 4 days (+tet/-bio) or with addition of both tetracycline for 4 days and biotin for 1 day (+tet/+bio); FLAG (green), Streptavidin (red), nuclei (blue). Scale bar: 20 µm. **D.** Principal component analysis (PCA) of the BioID data. Ellipses represent 95 % confidence intervals. **E.** Volcano plot of proteins enriched by streptavidin pull-down and analyzed by mass spectrometry from LMOD1-BirA* and LMOD2-BirA*. n = 4 biological replicates (Table S3) **F.** Quantification of selected unique interaction partners of LMOD1-BirA* and LMOD2-BirA* in comparison to BirA*-Ctrl. Each symbol represents a biological replicate, error bars indicate the SD. One-way ANOVA, *: p-value ≤ 0.05, **: p-value ≤ 0.01. **G.** Validation of the SIRT1-LMOD1 interaction. Cells expressing either GFP or SIRT1-GFP were used for co-immunoprecipitation using GFP-trap. The eluates from the GFP-trap were analyzed by immunoblot using antibodies directed to BirA* or GFP. For quantification, BirA* was normalized to SIRT1-GFP intensity. **H.** Representative images of proximity ligation assay (PLA) (red) in primary myoblasts during proliferation and after one day of differentiation. Scale bar: 10 µm. **I.** Representative immunofluorescence images of SIRT1 (gray), PAX7 (red), LAMININ (green), and Hoechst (blue) of uninjured TA section (day 0) and 5 days post-injury from young mice (3 months old).

**Figure S3:**
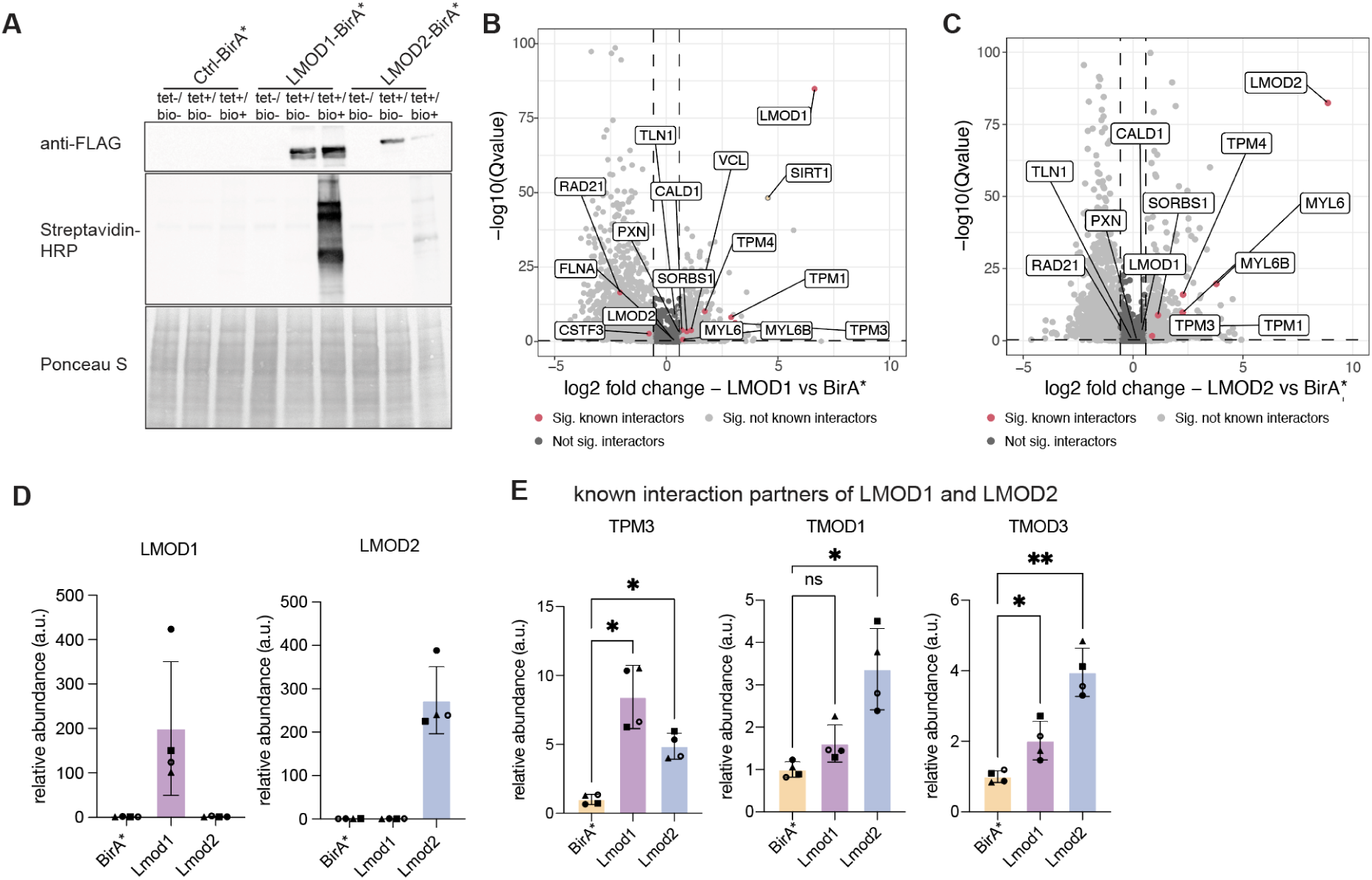
BioID for LMOD1 and LMOD2. **A.** Immunoblot of BirA* fusion proteins performed on lysates from HEK293T cells stably transfected with LMOD1-BirA*-FLAG, LMOD2-BirA*-FLAG or Ctrl-BirA*-FLAG following 24 h incubation with (+tet) or without (−tet) tetracycline. Middle panel, streptavidin-HRP blot following induction of BirA* fusion proteins with tetracycline and supplementation of biotin for 24 h. Ponceau stainings were used as loading control. HRP: horseradish peroxidase. **B**. Volcano plot of proteins enriched by streptavidin pull-down and analyzed by mass spectrometry from LMOD1-BirA* and **C.** LMOD2-BirA* compared to BirA*-Ctrl. Significantly enriched (AVG Log2 Ratio > 0.58 and Q value < 0.05), known interaction partners for LMOD1 and LMOD2 are highlighted in red, SIRT1 is highlighted in yellow (Table S3). **D.** Quantification of LMOD1 and LMOD2 protein abundance after overexpressing the LMOD1-BirA* and LMOD2-BirA* fusion proteins in comparison to BirA*-Ctrl. **E.** Quantification of selected known interaction partners of LMOD1-BirA* and LMOD2-BirA* in comparison to BirA*-Ctrl. For (**D**) and (**E**). Each symbol represents a biological replicate, and error bars indicate the SD. One-way ANOVA, *: p-value ≤ 0.05, **: p-value ≤ 0.01.

### LMOD1 and SIRT1 show dynamic subcellular localization during myogenic differentiation

Next, we investigated how LMOD1 might influence SIRT1 activity and thereby myogenic differentiation. SIRT1 has been shown to localize both in the cytoplasm and nucleus of neural precursor cells, murine pancreatic beta cells, rat cardiomyocytes, and vascular endothelial cells, while LMODs have been mainly associated with cytoplasmic localization (Hisahara et al. 2008; Tanno et al. 2010; Tong et al. 2013; Tanno et al. 2007; Conley 2001; Nauen et al. 2020). To determine the localization of SIRT1 and LMOD1 and a potential change in localization during myogenesis, we performed co-immunostainings for LMOD1 and SIRT1 in primary myoblasts to determine their localization at different stages of myogenic differentiation (Figure 5A). Overall, the total fluorescence intensity of LMOD1 significantly increased during differentiation (Figure S4A), while the signal intensity of SIRT1 slightly decreased (Figure S4B). However, under proliferating conditions, SIRT1 and LMOD1 predominantly localized in the nucleus (Figure 5A). Interestingly, after initiation of differentiation, both SIRT1 and LMOD1 were clearly detectable in the cytoplasm, indicating their translocation into the cytoplasm. At a later time point (2d of differentiation), the localization of SIRT1 can once again be observed in the nucleus, while LMOD1 remains in the cytoplasm. Using the fluorescent signal intensity, we quantified the cytoplasmic/nuclear ratios for both SIRT1 and LMOD1 and found that both proteins change their intracellular distribution during myogenic differentiation, suggesting that their localization in the cell affects the differentiation process (Figures 5B and 5C). Of note, we also observed a correlation between their cytoplasmic-nuclear partitioning, particularly during proliferation and at day one of differentiation (Figure 5D and 5E). The correlation between their respective localization was reduced at day 2 of differentiation, with SIRT1 displaying mainly a nuclear localization while LMOD1 remained predominantly cytoplasmic (Figure 5F). We confirmed these dynamic changes of sub-cellular localization for SIRT1 and LMOD1 by cellular fractionation experiments followed by immunoblot analysis (Figure S4C).

**Figure 5:**
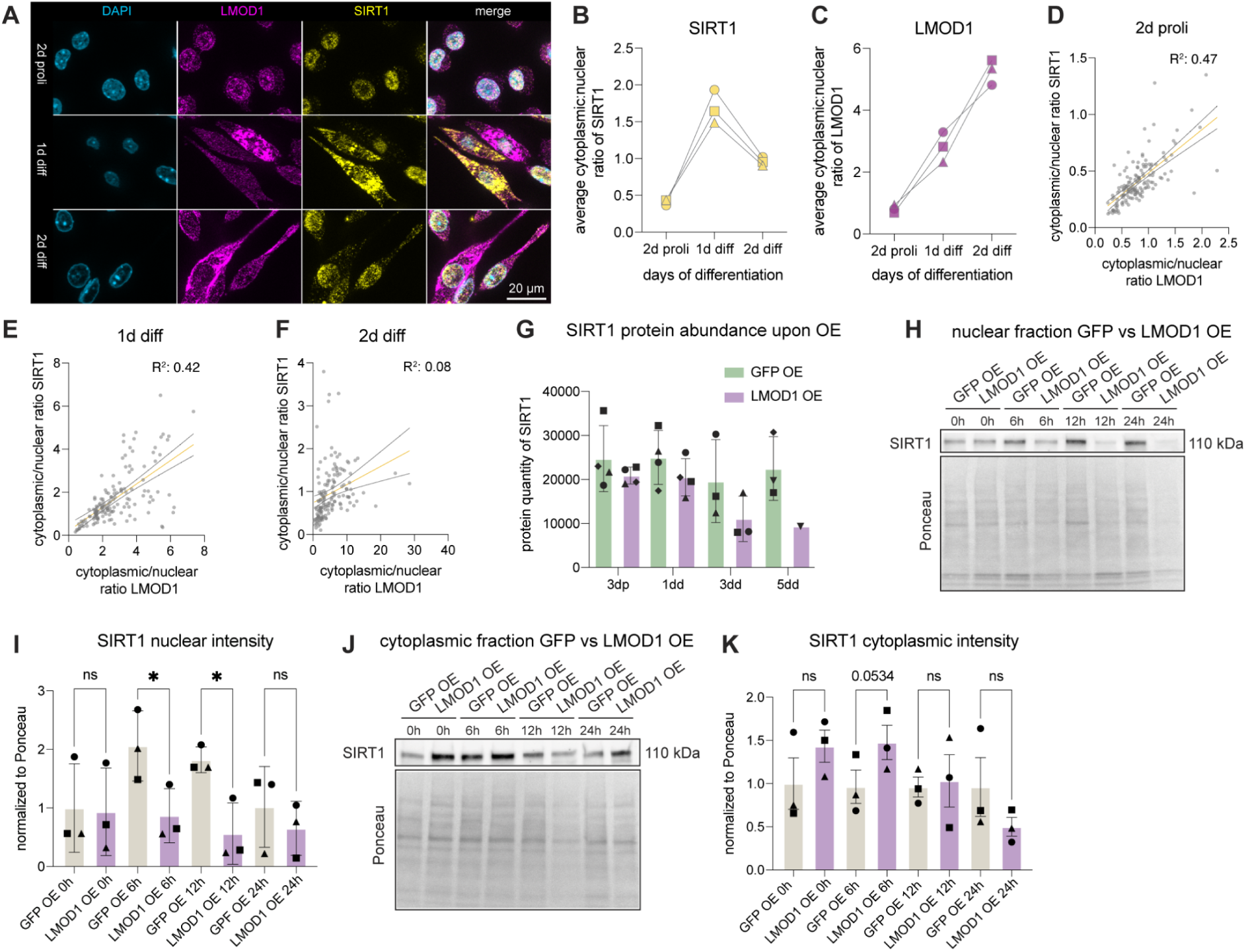
Overexpression of LMOD1 influences the subcellular localization of SIRT1. **A.** Representative immunofluorescence staining of LMOD1 and SIRT1 at different timepoints of myogenic differentiation; LMOD1 (purple), SIRT1 (yellow), nuclei (Hoechst, blue). Scale bar: 20 µm. **B - C.** Cytoplasmic to nuclear SIRT1 (**B**) and LMOD1 (**C**) ratio at different days of differentiation, n = 50 cells were analyzed per biological replicate per time point. **D - F.** Correlation of the cytoplasmic to nuclear ratio of LMOD1 and SIRT1 at 2 days of proliferation (R^2^ = 0.47) (**D**), 1 day differentiation (R^2^ = 0.42) (**E**) and 2 days of differentiation (R^2^ = 0.08) (**F**). **G.** SIRT1 protein abundance upon LMOD1 OE at different timepoints, 3 days proliferation (3dp), 1 day differentiation (1dd), 3 days differentiation (3dd), 5d differentiation (5dd). Each symbol represents a biological replicate (n = 4, primary myoblasts); error bars indicate SD. **H - K.** Representative immunoblot of SIRT1 in the nuclear (**H** and **I**) and cytoplasmic (**J** and **K**) fraction, comparing GFP and LMOD1 OE at early time points of differentiation (0h: undifferentiated/proliferating, 6h, 12h, 24h of differentiated primary myoblasts). SIRT1 signal was analyzed in n = 3 immunoblots and was normalized to the respective entire lane of the Ponceau staining. In bar plots, each symbol represents a biological replicate, and the error bars indicate the SD. Paired t-test *: p-value ≤ 0.05, ns: not significant.

Since SIRT1 protein abundance decreased with each day of differentiation as shown by our proteomics data and immunoblot analyses (Figure 5G and S4D), we hypothesized that the LMOD1-mediated change in SIRT1 localization is especially required during early differentiation. Therefore, we repeated the cellular fractionation experiments focusing on the onset of myogenic differentiation with additional time points and by using LMOD1 OE cells. Next, we compared the nuclear fraction of the LMOD1 OE cells to the control cells to determine if LMOD1 alone can alter SIRT1 subcellular localization (Figure 5H-K). In particular, at 6 and 12 hours of differentiation, SIRT1 is highly abundant in the nucleus in control cells. However, this dynamics shift and the SIRT1 abundance decreases in the nucleus when LMOD1 is overexpressed (Figure 5H and 5I). In line with this observation, in LMOD1 overexpressing cells the cytoplasmic SIRT1 abundance increased at 0h (proliferation) and 6h differentiation, suggesting a shift of SIRT1 protein to the cytoplasm upon LMOD1 OE compared to the control already under proliferating conditions (Figure 5J and 5K). Together, these data demonstrate that SIRT1 and LMOD1 dynamically change their subcellular localization and that overexpression of LMOD1 favors the cytoplasmic localization of SIRT1 at early stages of myogenic differentiation.

**Figure S4:**
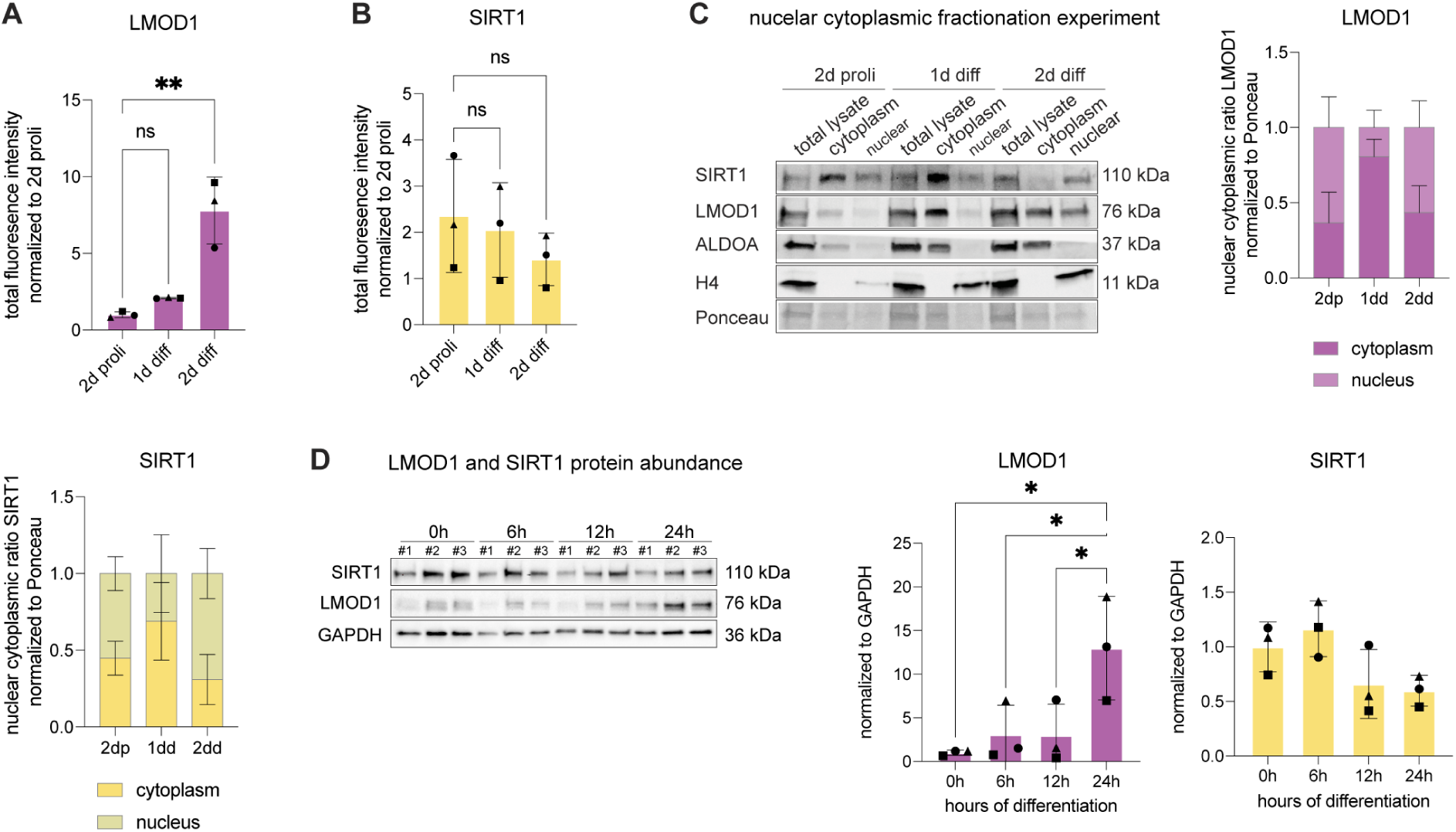
Sub-cellular localization of LMOD1 and SIRT1 during myogenic differentiation. **A - B.** Total immunofluorescence intensity (nucleus and cytoplasm) of LMOD1 (**A**) and SIRT1 (**B**) during different timepoints of myogenic differentiation. **C.** Representative immunoblot of the nuclear-cytoplasmic fractionation experiment in primary myoblasts at different timepoints: 2d proliferation, 1d differentiation and 2d differentiation. SIRT1 and LMOD1 signal was analyzed in (n = 3) immunoblots and normalized to the respective entire lane of the Ponceau staining. ALDOA signal was used as a cytoplasmic marker, and Histone 4 (H4) was used as a nuclear marker. Immunoblot analyses of LMOD1 and SIRT1 intensity for each cellular compartment are depicted in the barplots. One-way ANOVA *: p-value ≤ 0.05, ns: not significant. **D.** Immunoblot showing LMOD1 and SIRT1 protein abundance at different timepoints of myogenic differentiation (0h = proliferating conditions, 6h differentiation, 12h differentiation, 24h differentiation). Quantification of immunoblots for LMOD1 and SIRT1 shown in barplots normalized to GAPDH. For all bar plots, each symbol represents a biological replicate, and the error bars indicate the SD.

### Reduction or inhibition of SIRT1 can partially restore the impaired myogenic differentiation caused by knockdown of LMOD1

Given the interaction of LMOD1 and SIRT1, we hypothesized that LMOD1 regulates SIRT1-mediated transcription of genes that are critical for myogenic differentiation. To test this hypothesis, we investigated the impact of reduced SIRT1 levels using both a genetic (siRNA targeting *Sirt1*) and a pharmacological approach (25 µM EX527, small molecule inhibitor specific for SIRT1) during myogenic differentiation (Figure 6A) (Peck et al. 2010; Broussy et al. 2020). Primary myoblasts were transfected with siLmod1 and siSirt1 or incubated with EX527, at the time of induction of differentiation. After three days of differentiation, immunofluorescence staining was performed and analyzed (Figure 6B). We confirmed the significant decrease in LMOD1 and SIRT1 levels upon siRNA-mediated knockdown through immunoblot analysis (Figure S5A) and qRT-PCR (Figure S5B). Our results show that both genetic and pharmacological inhibition of SIRT1 signaling effectively promoted myogenic differentiation (Figure 6B and 6F) leading to an increased number of nuclei per myotube in the siLmod1/siSirt1 double knockdown compared to the knockdown of LMOD1 alone, indicating an improvement in myotube fusion after knockdown of SIRT1 (Figure 6C, 6G and Figure S5C and S5D). Moreover, both interventions were able to counteract the negative effects observed after knockdown of LMOD1 alone (Figure 6D and 6H). Previous studies have demonstrated that *Sirt1* knockout mice exhibited premature differentiation of MuSCs (Ryall et al. 2015). As a measure of myogenic differentiation, we measured the length of myotubes and observed an elongation after knocking down SIRT1 together with LMOD1 (Figure 6E and 6I).

**Figure 6:**
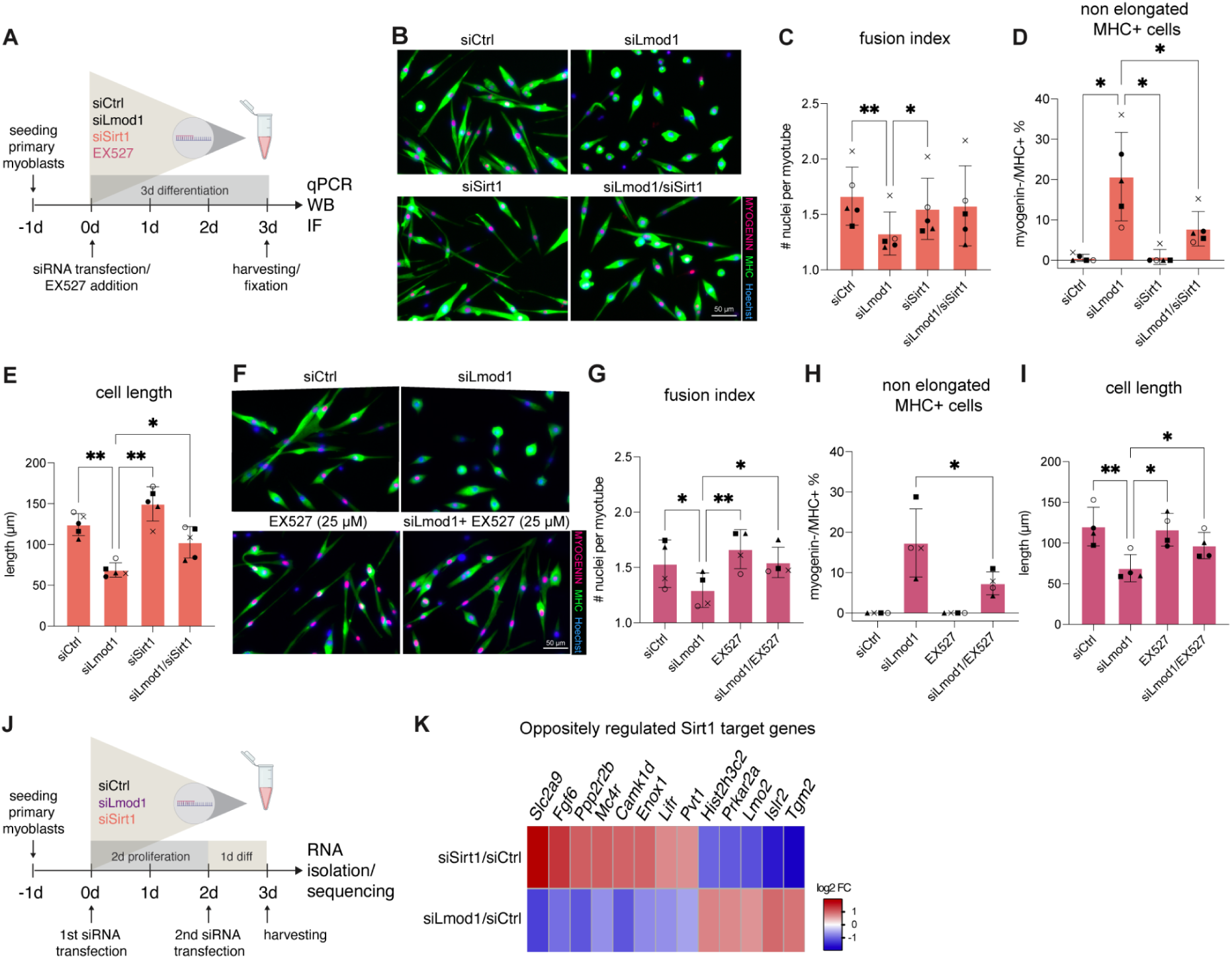
Reduced SIRT1 signaling can partially reverse siLmod1-induced impaired myogenic differentiation. **A.** Illustration of the experimental setup. Primary myoblasts were co-transfected with *Lmod1*-specific siRNA (siLmod1) and siRNA against Sirt1 (siSirt1) (depicted in orange) or treated with SIRT1 inhibitor EX527 (depicted in pink), simultaneously to the induction of differentiation. **B./F.** Representative immunofluorescence images after *Lmod1* and *Sirt1* knockdown with siRNA at day 3 of differentiation (**B**) or after *Lmod1* knockdown and addition of 25 µM EX527 at day 3 of differentiation (**F**); MYOGENIN (red), devMHC (green), nuclei (Hoechst, blue). **C - E.** Quantification of the number of nuclei per myotube (**C**), percentage of Myogenin-/MHC+ cells (**D**) and measured length of differentiated myotubes (**E**) after siRNA transfection of SIRT1 and LMOD1. One-way ANOVA *: p-value ≤ 0.05, **: p-value ≤ 0.01. **G - I.** Quantification of the number of nuclei per myotube (**G**), percentage of Myogenin-/MHC+ cells (**H**) and measured length of differentiated myotubes (**I**) after siRNA knockdown of LMOD1 and EX527 treatment to inhibit SIRT1. One-way ANOVA *: p-value ≤ 0.05, **: p-value ≤ 0.01. **J.** Illustration of the experimental setup for the RNA-Seq experiment. Primary myoblasts were first seeded and then transfected with siRNA against LMOD1 or SIRT1 or siCtrl. After two days of incubation, the primary myoblasts were transfected a second time with siRNA and induced to differentiate simultaneously. After one day of differentiation, cells were harvested, and RNA was isolated for library preparation and RNAseq analysis. **K.** Heatmap indicating oppositely differentially expressed SIRT1 target genes from siSirt1 vs siCtrl an siLmod1 vs siCtrl with a log2 FC ratio > 0.58. SIRT1 target genes identified from ChipSeq experiment published in (Table S4) (Ryall et al. 2015). For all bar plots, each symbol represents a biological replicate, and the error bars indicate the SD.

Next, we performed RNA-sequencing to identify downstream effectors that could explain the improvements in myogenic differentiation observed after double knockdown of *Lmod1* and *Sirt1* (Table S4). Therefore, we performed a knockdown of *Lmod1* and *Sirt1* two days prior to differentiation, followed by a second siRNA transfection at the initiation of differentiation to ensure a high knockdown efficiency. After one day of differentiation, cells were harvested to quantify early transcriptional changes in the myogenic program after knockdown of *Lmod1* and *Sirt1* using RNAseq (Figure 6J). Differential gene expression analysis confirmed a significant reduction of *Lmod1* and *Sirt1* mRNA in our experimental setup (Figure S5E). Next, we identified 246 genes being significantly and oppositely regulated after knockdown of either *Lmod1* or *Sirt1* (Table S4). Interestingly, among these genes, we identified a subset of known SIRT1 target genes that were previously defined by ChIP-Seq analysis of MuSCs from WT and Sirt1^mKO^ mice (Ryall et al. 2015) (Table S4). These include key factors involved in cell signaling and differentiation, such as *Fgf6*, *Tgfa*, *Mcr*, *Lifr*, and *Camk1d* (Figure 6K). Taken together, our data show that genetic or pharmacological inhibition of SIRT1 signaling can partially reverse the impaired myogenic differentiation caused by knockdown of *Lmod1* and that affecting LMOD1 levels is sufficient to alter the expression of a subset of SIRT1 target genes, highlighting the role of the LMOD1-SIRT1 axis in myogenic differentiation.

**Figure S5:**
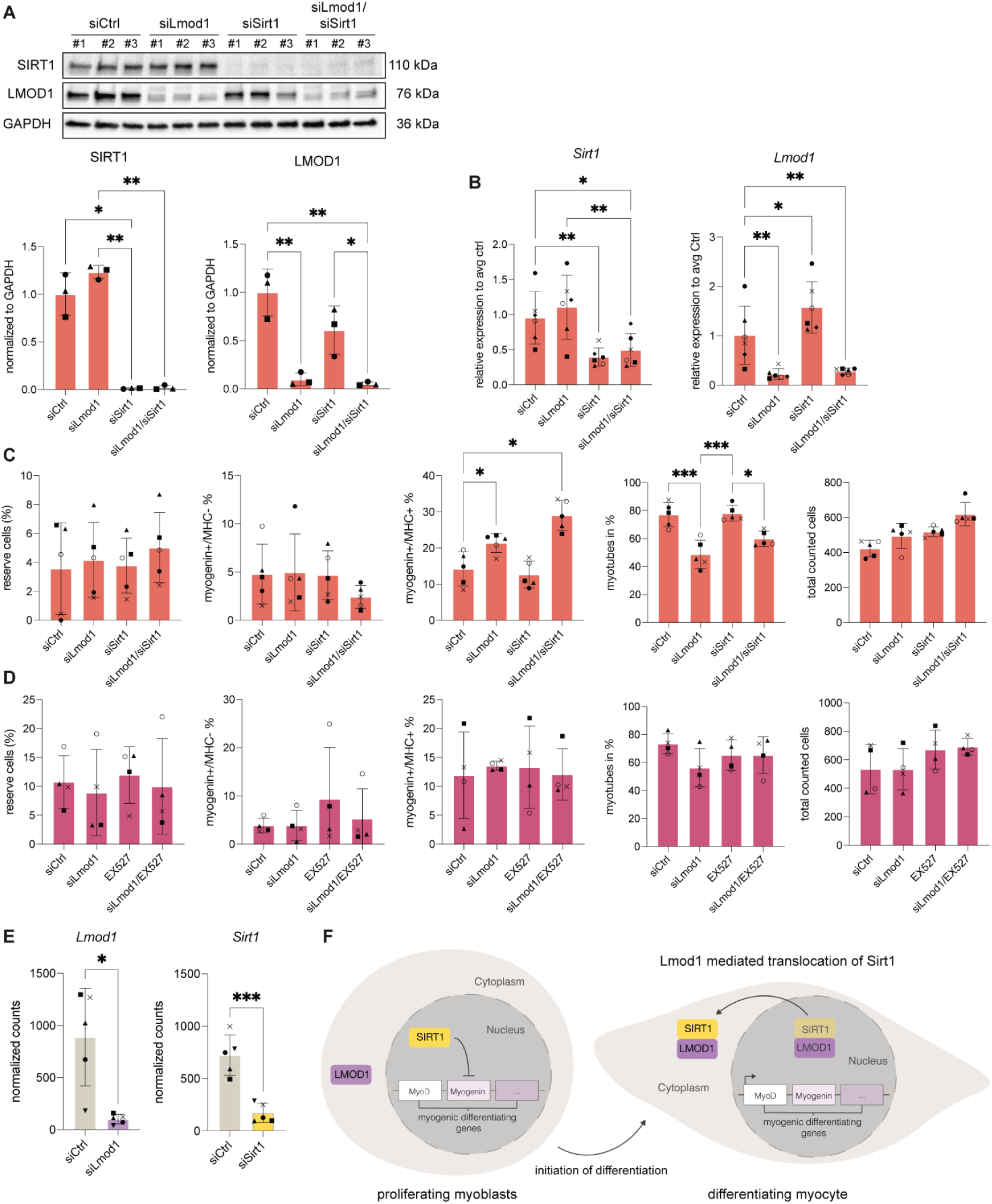
Reduced SIRT1 signaling can partially reverse siLmod1-induced impaired myogenic differentiation. **A.** Immunoblot validation of the LMOD1 and SIRT1 single knockdown and double knockdown (siLmod1/siSirt1) or a scrambled siRNA. Primary myoblasts were transfected at the initiation of differentiation and harvested after three days of differentiation. One-way ANOVA: *: p-value ≤ 0.05. **: p-value ≤ 0.01. **B.** qRT-PCR analysis showing the relative expression of *Sirt1* and *Lmod1* in siLmod1 or siSirt1 single knockdown or double knockdown compared to siCtrl treated cells, normalized to *Gapdh* expression levels. Paired t-test *: p-value ≤ 0.05 **: p-value ≤ 0.01. **C** and **D.** Percentage of cell populations found in immunofluorescence staining: non-proliferating myoblasts (Hoechst+/Myogenin-/MHC-) cells, Myogenin+/MHC-cells, Myogenin+/MHC+, fully differentiated myotubes and total counted cells under differentiating conditions after siRNA mediated knockdown of LMOD1 and SIRT1 (**C.** related to Figure 6C - E) or siRNA mediated knockdown of LMOD1 and EX527 treatment to inhibit SIRT1 (**D.** related to Figure 6G - I.) **E.** Normalized RNA-seq read counts of *Lmod1* and *Sirt1* after one day of differentiation after siCtrl and siLmod1 and siCtrl and siSirt1 knockdown (Table S4). Paired t-test *: p-value ≤ 0.05 ***: p-value ≤ 0.001. For all bar plots, each symbol represents a biological replicate, and the error bars indicate the SD. **F.** Working model: During the proliferation stage, SIRT1 is found in the nucleus, repressing the expression of myogenic regulating factors. However, at the onset of myogenic differentiation, LMOD1 interacts with SIRT1 and affects its subcellular localization, leading to decreased levels of SIRT1 in the nucleus. This reduction in SIRT1 levels results in the de-repression of MRFs and the initiation of differentiation.

## Discussion

Myogenic differentiation involves the coordinated alterations of gene expression and reorganization of the cellular architecture. Here, we present a temporally resolved analysis of the proteome of primary mouse myoblasts undergoing differentiation *in vitro.* With a depth of more than 6000 proteins, our analysis covers three times more proteins compared to a previous study performed on an immortalized myoblast cell line (C2C12 cells) (Tannu et al. 2004; Kislinger et al. 2005). By using five independent primary myoblast lines, we were able to take into account the variability that typically affects primary cultures and derive robust and reproducible proteome signatures (Hindi et al. 2017; Kim et al. 2020). Our data highlight hundreds of changes in energy metabolism, RNA and protein synthesis, as well as cytoskeletal organization that occur with specific temporal dynamics during myogenic differentiation. To demonstrate the relevance of our data, we decided to focus on LMOD1, a cytoskeletal protein whose role in myogenic differentiation has not yet been investigated. Although it is known that actin cytoskeletal dynamics are involved in myogenesis and that LMOD1 contributes to actin filament nucleation in smooth muscle cells (Guerin and Kramer 2009; Heng and Koh 2010; Xie et al. 2020; Chereau et al. 2008), the biological significance and functional role of LMOD1 during myogenic differentiation have remained elusive so far. We found that LMOD1 is expressed by MuSCs, that its expression increases during myogenic differentiation *in vitro* and *in vivo*, and that reduction of LMOD1 levels significantly affects myotube formation and the number of myonuclei per myotube. Furthermore, the timing of induction of LMOD1 expression during skeletal muscle regeneration suggests a role in differentiation and fusion rather than in myofiber maturation. This is further supported by our finding that knockdown of *Lmod*1 after injury of skeletal muscle results in reduced regeneration. Consistently, our knockdown studies revealed that the reduction of LMOD1 severely affects the myogenic differentiation and myonuclear numbers. To confirm that these effects were indeed derived from a specific functional disruption rather than poor cell viability, we performed a TUNEL assay, which showed no increase in apoptosis upon LMOD1 knockdown. This suggests a model where the observed phenotype reflects an arrested differentiation program, rather than LMOD1 being required for cell survival. This is further supported by our finding that knockdown of LMOD1 after skeletal muscle injury resulted in impaired regeneration.

Cells that expressed Myogenin started differentiating into proper myotubes, indicated by the quantification of Myogenin+/MHC- and Myogenin+/MHC+ cells even after knockdown of LMOD1. However, a subset of cells positive for MHC but negative for Myogenin did not progress in differentiation, possibly due to loss or insufficient activation of myogenin expression. Loss of myogenin has been shown to lead to deregulated mTORC1 signaling and results in faster cell cycle entry in MuSCs (Ganassi et al. 2020). Since the knockdown of LMOD1 leads to an increase in the percentage of Myogenin-/MHC+ cells, the observed differentiation phenotype might be at least partially explained by a direct or indirect effect of LMOD1 on myogenin expression.

We found that LMOD1 selectively interacts with SIRT1. So far, there is no direct evidence suggesting that LMOD1 and SIRT1 have a specific binding domain where they can directly bind to each other. However, the comparison of primary sequences between LMOD1 and LMOD2 (Fowler and Dominguez 2017) reveals a 103 amino acid sequence unique to LMOD1, which could represent a possible binding interface. It is also conceivable that other proteins or molecules might be involved in mediating the interaction between LMOD1 and SIRT1, e.g., the RuvB-like AAA ATPase 1/2 (RUVBL1/RUVBL2). RUVBL1 participates in various complexes, including INO80, NuA4, SWR1, TIP60-P400, PAQosome (R2TP), as a transcriptional and/or chromatin modifier, with or without its homologue RUVBL2 (Gorynia et al. 2011; Chen et al. 2015; Lakisic et al. 2016).

SIRT1 is a protein deacetylase that can localize in the nucleus and cytoplasm (Bai and Zhang 2016; Moynihan et al. 2005; Chen et al. 2006), and was shown to play a critical role in muscle differentiation and metabolism (Ryall et al. 2015; Diaz-Ruiz et al. 2015; Fulco et al. 2003, 2008). Myogenesis involves the shuttling of various proteins and myogenic regulators between the nucleus and cytoplasm, regulated by various signals and interactions (Grifone et al. 2021; Nguyen et al. 2022). We have observed that the localization of SIRT1 changes dynamically during myogenic differentiation, and that the interaction between SIRT1 and LMOD1 increases during the initial stages of myogenic differentiation. Additionally, we showed that overexpression of LMOD1 affects the subcellular localization of SIRT1, leading to reduced levels of SIRT1 in the nucleus. We hypothesize that relocation of SIRT1 to the cytoplasm, mediated by LMOD1, is critical for the initiation of myogenic differentiation, resulting in the de-repression of MRFs, including myogenin. Alternatively, LMOD1 might sequester SIRT1 within the cytoplasm, preventing its re-entry into the nucleus, which in turn results in the expression of MRF to initiate myogenic differentiation (Figure S5F).

Our immunostaining data show that while LMOD1 is consistently cytoplasmic, its partner SIRT1 is only transiently localized in the cytoplasm. This suggests that their interaction is dynamically regulated. We hypothesize that the function of LMOD1 is determined by the changing availability of its binding partners during differentiation. During the initial phase, LMOD1 may primarily function to sequester SIRT1, a key regulator of myogenic genes. As differentiation proceeds, the increased expression of cytoskeletal components, such as its canonical partners TMODs and TPMs, likely shifts the function of LMOD1 towards its role in actin nucleation. This molecular switch, potentially driven by a change in the interactome of LMOD1, could then result in the release of SIRT1 from the cytoplasm. Such a mechanism may coordinate transcriptional regulation with cytoskeletal remodeling during myoblast differentiation.

Previous studies have demonstrated that reducing the protein levels of SIRT1 can enhance the differentiation of MuSCs (Ryall et al. 2015). Consistently, our data suggest that the negative impact of knockdown of *Lmod1* on myogenic differentiation can be partially restored by co-depletion of *Sirt1*. Our RNAseq analysis highlighted a subset of genes that were oppositely up- or downregulated after knockdown of either LMOD1 or SIRT1, including secreted factors and receptors that could potentially mediate this restoration of myogenic differentiation when LMOD1 and SIRT1 levels are reduced. The identified factors include *Fgf6*, *Tgfa*, *Camk1d*, *Mcr*, *Lifr*, *Isl2* and *Tgm2,* which were previously shown to be involved in proliferation and differentiation of MuSCs (Armand et al. 2006; Knight and Kothary 2011; Yamane et al. 1998; Dos Santos et al. 2023; Hunt et al. 2013) by modulating signaling pathways such as mTOR, Wnt or IGF1-PI3K/Akt-Foxo signaling (Wang et al. 2022; Cui et al. 2020; Zhang et al. 2018). Further investigation of possible downstream effectors or modulators identified by RNAseq could reveal additional mechanisms by which LMOD1 interferes with the myogenic program. Moreover, delineating the functional specialization and potential redundancy among leiomodin proteins represents an important next step. Our data indicate that LMOD1 primarily regulates early myogenic differentiation (Figure 3). In contrast, the lack of an early functional phenotype upon LMOD2 depletion, together with its upregulation at later stages (Figure S2A), suggests a temporal shift in regulatory control. Accordingly, a systematic comparative analysis of LMOD1, LMOD2, and LMOD3 will be required to elucidate their distinct roles in actin cytoskeleton regulation across the myogenic program, particularly with respect to myofibril maturation and maintenance.

Finally, we have discovered that the abundance of LMOD1 is approximately four times higher in MuSCs from geriatric mice compared to the abundance in MuSCs from young mice. Specifically, LMOD1 tends to accumulate in the nucleus of MuSCs as mice age. Disruption of the actin organization of the cytoskeleton and dynamics in aging and disease, such as Duchenne muscular dystrophy (DMD) or Nemaline myopathy (NM) was shown to impact the expression of myogenic regulators and different signaling pathways (Lai and Wong 2020; Eliazer et al. 2019; Kim et al. 2022). Studies have shown that SIRT1 maintains H4K16 in a deacetylated state in quiescent MuSCs, which leads to transcriptional repression (Ryall et al. 2015). Given the SIRT1-LMOD1 interaction that we have discovered, we speculate that the nuclear accumulation of LMOD1 might affect gene expression and chromatin remodeling in aged MuSCs. However, it remains to be investigated whether the accumulation of LMOD1 is a compensatory mechanism or is a causative factor leading to the dysregulation and impairment of MuSCs during aging. Identifying the molecular mechanism and potential downstream targets could be a promising strategy for discovering potential therapeutic targets for promoting muscle regeneration and improving muscle health in older adults.

## Acknowledgments

The authors gratefully acknowledge support from the FLI Core Facilities Proteomics, Sequencing, Imaging, and the Mouse Facility. A.O. is supported by the German Research Council (Deutsche Forschungsgemeinschaft, DFG) via the Research Training Group ProMoAge (GRK 2155), the Else Kröner Fresenius Stiftung (award number: 2019_A79), the Fritz-Thyssen Foundation (award number: 10.20.1.022MN), the Chan Zuckerberg Initiative Neurodegeneration Challenge Network (award numbers: 2020-221617, 2021-230967 and 2022-250618), the NCL Stiftung, and the Leibniz Association (XpandHSC, project ID K243/2019). J.vM. received funding from the Deutsche Forschungsgemeinschaft (MA-3975/2-1), the Deutsche Krebshilfe and the Carl Zeiss Foundation (DKH-JvM-861005). S.C.S. was supported by a postdoctoral fellowship by the German Research Foundation (DFG, 505064275). The FLI is a member of the Leibniz Association and is financially supported by the Federal Government of Germany and the State of Thuringia.

## Author contributions

Conceptualization: ES, SCS, JvM, AO Data curation: ES, SCS, AO Investigation: ES, IH, MH, AM, JvM, KH Methodology: ES, SCS, TD, JvM Project administration: JvM, AO Data analysis: ES, MH, AO Supervision: JvM, AO Visualization: ES, AO Writing – original draft: ES, JvM, AO Writing – review & editing: SCS

## Declaration of interest

Authors declare no competing interests.

## EXPERIMENTAL MODEL AND SUBJECT DETAILS

### Mice

All wild-type mice were C57BL/6J obtained from Janvier or internal breeding at the Leibniz Institute on Aging – Fritz Lipmann Institute (FLI) using the Janvier strain. All animals were kept in groups in a specific pathogen-free animal facility with a constant 12 h light/dark cycle and fed *ad libitum*. Young mice were aged 2-6 months, old mice were aged 18-20 months and geriatric animals were aged 24-33 months. All animal experiments were approved by the Thüringer Landesamt für Verbraucherschutz (Germany) under Reg.-Nr. FLI-17-014 and FLI-22-011.

### Cardiotoxin muscle injury and siRNA injection of the TA muscle

Young mice (3-months old) were anaesthetized by inhaling isoflurane using the Tec 7 Isoflurane (798932, Covetrus) Anesthesia Vaporizer (Datex Ohmeda). The TA muscle was injured once by injecting 50 μl of 10 μM cardiotoxin (L8102, Latoxan) in sterile 0.9% NaCl (12391112, Medpex) using an insulin syringe with a 29 G needle. To manage pain, the mice were given Metacam (1 mg/kg, 798932, Covetrus) subcutaneously for three days.

Self-delivering Accell siRNA ON-TARGETplus Non-targeting Pool (D-001910-10-50, Dharmacon) or Accell siRNA ON-TARGETplus targeting *Lmod1* (E-051082-00-0050, Dharmacon) were injected into the CTX-damaged TA-muscles on day 3 after injury to induce knockdown *in vivo*. TA muscles were further analyzed by quantifying the number of myogenin+ cells per area.

### FACS isolation of MuSCs

MuSCs were isolated from the hindlimb muscles of adult male mice of age 8 - 12 weeks. The muscles were dissected and collected in PBS, minced using scissors and digested in 2.5 g/mL collagenase B (Sigma) and 1 g/mL dispase (Sigma) for 30 min at 37 °C. Together with 12 mL of isolation medium (HAMs F10 + 20 % FBS) the digested muscle pieces were transferred into a 15 mL tube to allow the sedimentation of larger undigested muscle chunks. The sample was then filtered using a 0.74 mm cell strainer into a 50 mL tube, spun down at 450x *g* for 5 min at room temperature and resuspended in 500 mL isolation medium. The sample was then incubated on ice for 15 min with the antibodies used for FACS analysis, as indicated below. Afterwards, 15 mL PBS was added, and the sample was spun down for 5 min at 450x *g* at room temperature followed by resuspension in 1mL PBS. After filtering the cells with a 0.35 mm cell strainer staining with SYTOX Blue Dead Cell Stain (S34857, Thermo Fisher) was performed to distinguish between dead and alive cells. A BD FACSAria III was used for sorting the cells. To FAC sort MuSCs, gating was set to alpha7-integrin+ (anti-Alpha 7 Integrin 647 (clone:R2F2), 67-0010-05, The University Of British Columbia AbLab, 1:500), CD11b-(CD11b-PE, 553311, BD Bioscience, 1:500), CD31-(CD31-PE, 553373, BD Bioscience,1:500), CD45-(CD45-PE, 553081, BD Bioscience,1:500) and Sca1-(Ly-6A/E-PE, 553108, BD Bioscience, 1:500).

### Cell culture

FACS isolated primary myoblasts were cultured on collagen (#354236, Corning) coated (1h at RT with 0.167 mg/mL) plates in Ham’s F-10 Nutrient Mix (Thermo Fisher #31550031) supplemented with 10 % FBS (#10270106, Thermo Fisher), 1 % Penicillin-Streptomycin (#15140-122, Thermo Fisher) and 2.5 ng/mL bFGF (#13256029, Thermo Fisher) in a 37 °C incubator with 95 % humidity and 5 % CO_2_. For the proliferation assay, primary myoblasts were seeded onto collagen-coated 24-well plates (734-2325, VWR) (25,000 cells per well) and cultured for 48 h in the myoblast growth medium. For differentiation assays on 24-well plates, 75,000 primary myoblasts were seeded and cultivated for 72 h in the differentiation medium. Primary myoblasts were always harvested using 0.25 % Trypsin. For location stainings, cells were grown on collagen-coated 4-well culture slides (80426, Ibidi). For proliferation conditions, 5,000 cells were seeded per well, and 12,000 cells were seeded per well for differentiation conditions. The cells were then incubated at 37 °C. HEK293T cells (Flp-In 293 T-Rex cells) were obtained from Thermo Fisher (R78007). HEK293T cells were grown in Dulbecco’s modified Eagle’s medium (#D6429, Sigma) with high glucose (5 g/l) supplemented with 10 % heat-inactivated FBS and supplementation with 100 mg/mL Zeocin (#R250-01, Thermo Fisher) and 15 mg/mL Blasticidin (#R210-01, Thermo Fisher). After generation of a stable line cells were supplemented with 100 mg/mL Hygromycin (#10687-010, Thermo Fisher) and 15 mg/mL Blasticidin. Cells were cultured in a 37 °C incubator with 95 % humidity and 5 % CO_2_. For immunofluorescence staining of HEK293T, cells were grown on 50 μg/mL Poly-D-Lysine solution (Gibco™ A3890401) coated autoclaved coverslips (Carl Roth, YX03.1). Each coverslip was placed in an individual well of a 12-well Lab solute plate (with a specified number), and 25,000 cells were seeded per well. The Platinum-E (Plat-E) cells, a potent retrovirus packaging cell line were cultured in DMEM high glucose (#D6429, Sigma) supplemented with 10 % FBS, 1 %Penicillin-Streptomycin in a 37 °C incubator with 95 % humidity and 5 % CO_2_.

### siRNA transfection of cells with Lipofectamine® RNAiMAX Reagent

Knockdown experiments using small interfering RNAs (siRNA) (ON-TARGETplus Non-targeting Pool (D-001810-10-05, Dharmacon), SMART:pool ONTARGETplus Lmod1 siRNA (9530015K06Rik | SM-Lmod), SMART:pool ONTARGETplus Lmod2 siRNA (C-Lmod), SMART:pool ONTARGETplus Sirt1 siRNA (AA673258 | SIR2L1 | Sir2 | Sir2a | Sir2alpha) against a target mRNA were performed. For siRNA transfection, the Lipofectamine® RNAiMAX Kit (13778075, ThermoFisher) was used according to the manufacturer’s protocol.

### Generation of overexpression cell lines

The murine stem cell virus (MSCV) retroviral expression system (634401, Takarabio) was used for overexpression studies. The MSCV vectors are optimized for introducing and expressing target genes in stem cells such as MuSCs. The vector pENTR233.1-Lmod1 (100016603, Dharmacon, spectinomycin resistant) and GFP were blunt end cloned in the donor vector cloneJET^TM^ (K1232, Thermo Fisher, ampicillin-resistant) to finally generate pMSCV.puro plasmids (634401, Takarabio, ampicillin-resistant) encoding Lmod1 and GFP, respectively (Table 1).

**Table 1:**
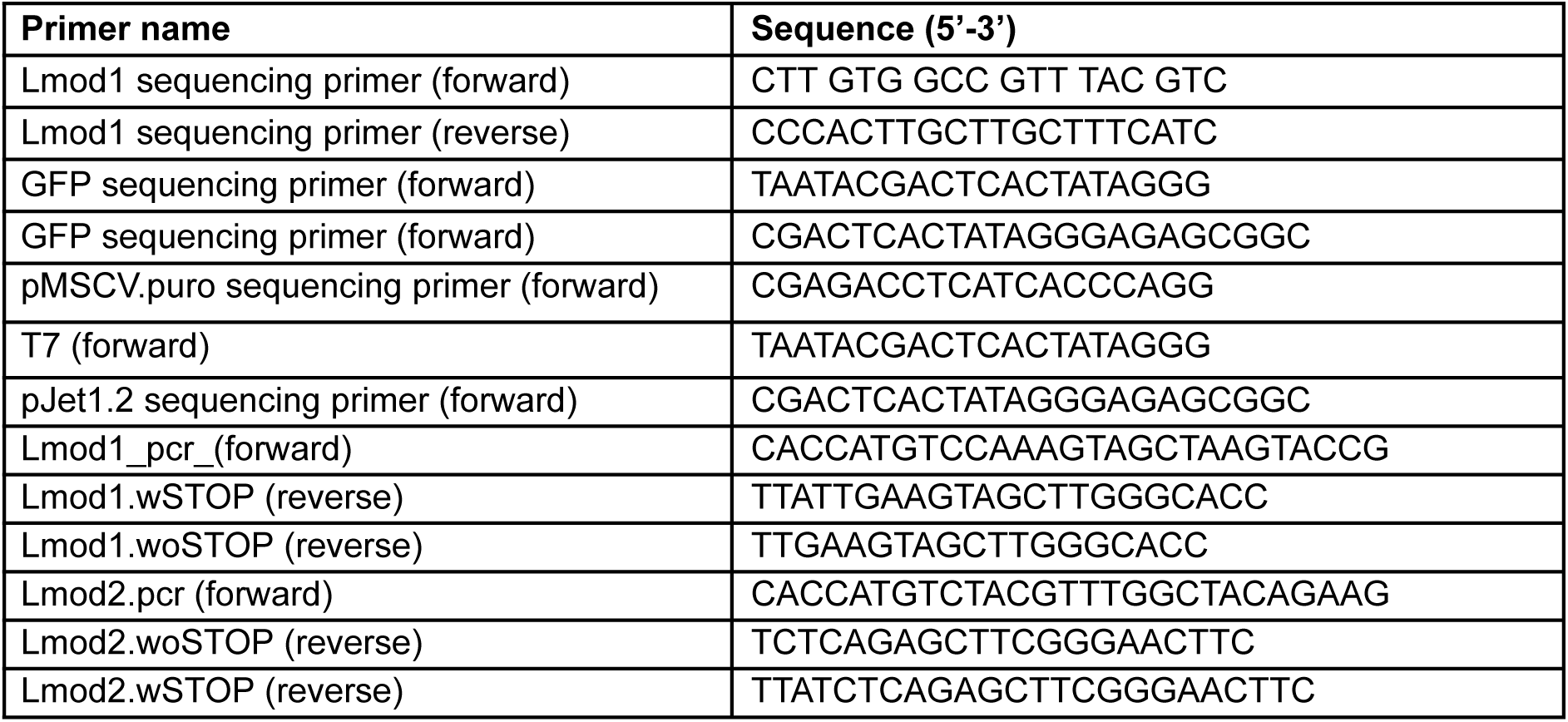
Primers for cloning.

Plat-E cells were transfected with 20 µg of the generated pMSCV.puro plasmids using polyethylenimine (PEI). 60 uL PEI (1 mg/mL) were added to the plasmid mixture (3:1 ratio of PEI:DNA), briefly mixed by inverting the tube and incubated for 20 min at RT. Subsequently, the transfection complex was added dropwise to a 10 cm plate and incubated for 5-6 h in 3.5 % CO2, 37 °C. The medium was changed to standard proliferation medium (DMEM, 10 % FBS and 1 % Pen-Strep) and incubated until the next day to collect the retrovirus. For further infection of primary myoblasts, 1 Mio cells were seeded in a p10 plate and infected using a 1:5 ratio of retrovirus to growth medium (F10, 20 % FBS, bFGF, 1 % Pen-Strep) mixed with 5 µl of Protamine Sulfate (8 µg/mL). Infection of primary myoblasts was repeated the next day as previously described and incubated at 37 °C. After 48 h post-transfection, primary myoblasts were selected with 1.25 µg/mL puromycin for up to one week. Medium was changed every 2 - 3 days.

### *In vitro* proximity labeling for BioID protocol

HEK293T cells (Flp-In 293 T-Rex cells) expressing BirA*-LMOD1 and BirA*-LMOD2, were generated as described by (Mackmull et al. 2017). The cells were selected using 15 μg/mL Blasticidin (GibcoTM, R21001) and 100 μg/mL Zeocin™ (Thermo Fisher Scientific, R25001). Once stable cell lines were generated, the cells were selected using 100 μg/mL Hygromycin B (Thermo Fisher Scientific, 10687010) and 15 μg/mL Blasticidin (GibcoTM, R21001).

Stable HEK293T cell lines expressing fusion proteins containing BirA* were seeded in 150 mm dishes at the density of approximately 500K cells. To induce expression of BirA* fusion proteins, cells were treated with 1 μg/mL tetracycline (Sigma Aldrich, 87128) for four days. Biotin (50 μM) (Sigma Aldrich, B4501) was added to the cells 24 h prior to harvest to biotinylate proximal proteins.

### Skeletal muscle cryostat sections and staining

TA muscles were isolated and frozen sections (14 μm) were made using a Leica Cryostat CM 3050 (Meyer Instruments) at −21 °C and placed on glass slides. Sections were stored at −80 °C. For immunostaining, muscle cryosections were fixed with 4 % paraformaldehyde (v/v, CP10.1, Roth, pH 7.4). for 7 min and then permeabilized with 0.1 M Glycine (1042011000, VWR) in PBS, (pH 7.4) for 5 min and after one wash with PBS 0.5 %Triton-X was added (v/v, 3051.3, Roth) for 10 min. After another washing step with PBS sections were incubated for 20 min at 65 °C in pre-heated Citrate Antigen Retrieval buffer (10 mM Sodium citrate, 0.05 % Tween 20, pH 6.0). Slides were re-equilibrated by washing 3x with PBS for 5 min and incubated for 30 min using MOM blocking (1:40) and subsequently blocked in blocking solution (5 % horse serum (v/v) in PBS) at RT. The sections were incubated in primary antibody Pax7 (Pax7, DSHB, Hybridoma mouse IgG1, undiluted), myogenin (DSHB, F5D, hybridoma cell supernatant, undiluted), Lmod1 (Proteintech, 15117-1-AP, 1:500) and Laminin (LSBio, LS-C96142, 1:500)) overnight at 4 °C in a wet chamber. Sections were washed 3x with 0.1 % Triton in PBS (pH 7.4) for 5 min and secondary antibodies anti-chicken IgG (Alexa Fluor® 488, A-11039), anti-mouse IgG (Alexa Fluor® 546, A-21123, Thermo Fisher, 1:1000) and anti-rabbit IgG (Alexa Fluor® 647, A-31573, Thermo Fisher, 1:1000), were applied for 1 h at RT. Nuclei were stained with Hoechst (bisBenzimideH 33258, B2261 Sigma, 0.02 μg/μl in PBS, 1:5000) for 5 min. After washing sections 3x with 0.1 % Triton in PBS, sections were mounted in Permafluor mountain medium (TA-006-FM, Thermo Fisher). Immunofluorescence microscopy was performed with the Zeiss Axio Scan Z.1 using a Plan-Apochromat 20x/0.8 M27 Objective (Zeiss).

### Immunofluorescence staining of primary myoblasts

Primary myoblasts from proliferation or differentiation assays were fixed with 2 % formaldehyde (v/v in PBS, CP10.1, Roth) for 5 min at RT, washed twice with PBS for 5 min and were permeabilized with permeabilization solution (0.1 % Triton-X-100 (v/v, 3051.3, Roth), 0.1 M Glycine (1042011000, VWR) in PBS) for 5 min at RT. Cells were incubated for 1 h in blocking solution (5 % horse serum, v/v, 26050-088, Thermo Fisher) in PBS at RT. Incubation with primary antibodies was carried out in 5 % horse serum in PBS overnight at 4 °C. Subsequently, the cells were washed three times with PBS for 5 min and incubated with the secondary antibodies in blocking solution for 1 h at RT in the dark. Next, the cells were washed once with PBS for 5 min, stained with Hoechst (bisBenzimide H 33258, B2261 Sigma, 1:5000 in PBS) at RT for 5 min and washed twice with PBS for 5 min. The cells were stored at 4 °C in PBS in the dark until they were analyzed. For location stainings, the removable chamber was detached, and cells were mounted in permafluor mounting medium (TA-006-FM, Thermo Fisher) and coverslips (631-1574, VWR). The cells were then analyzed using an Axio Observer microscope (Carl Zeiss) or with an Axio Imager (Z2 using a Plan-Apochromat 63 x / 0.8 M27 Objective) and analyzed with Zen 2 Blue Edition software (Carl Zeiss Microscopy GmbH).

Primary antibodies used for immunofluorescence analysis (Table 3): Hybridoma mouse IgG1 myogenin (DSHB, F5D, hybridoma cell supernatant, undiluted), Hybridoma mouse IgG1 devMHC (DSHB, MF20, hybridoma cell supernatant, undiluted), anti-ki67 (ab15580, rabbit IgG 1:500), Lmod1 (Proteintech, 15117-1-A, rabbit, IgG, 1:500) and Sirt1 (Proteintech, 60303-1-Ig, mouse, IgG2b, 1:400). Secondary antibodies: anti-mouse IgG2b (Alexa Fluor® 488, A-21141, Thermo Fisher, 1:1000), anti-rabbit IgG (Alexa Fluor® 546, A10040, Thermo Fisher, 1:1000), anti-mouse IgG (Alexa Fluor® 546, A-21123, Thermo Fisher, 1:1000). Tunel staining was performed using the In Situ Cell Death Detection Kit (Roche) following the instructions from the manufacturer.

**Table 2:**
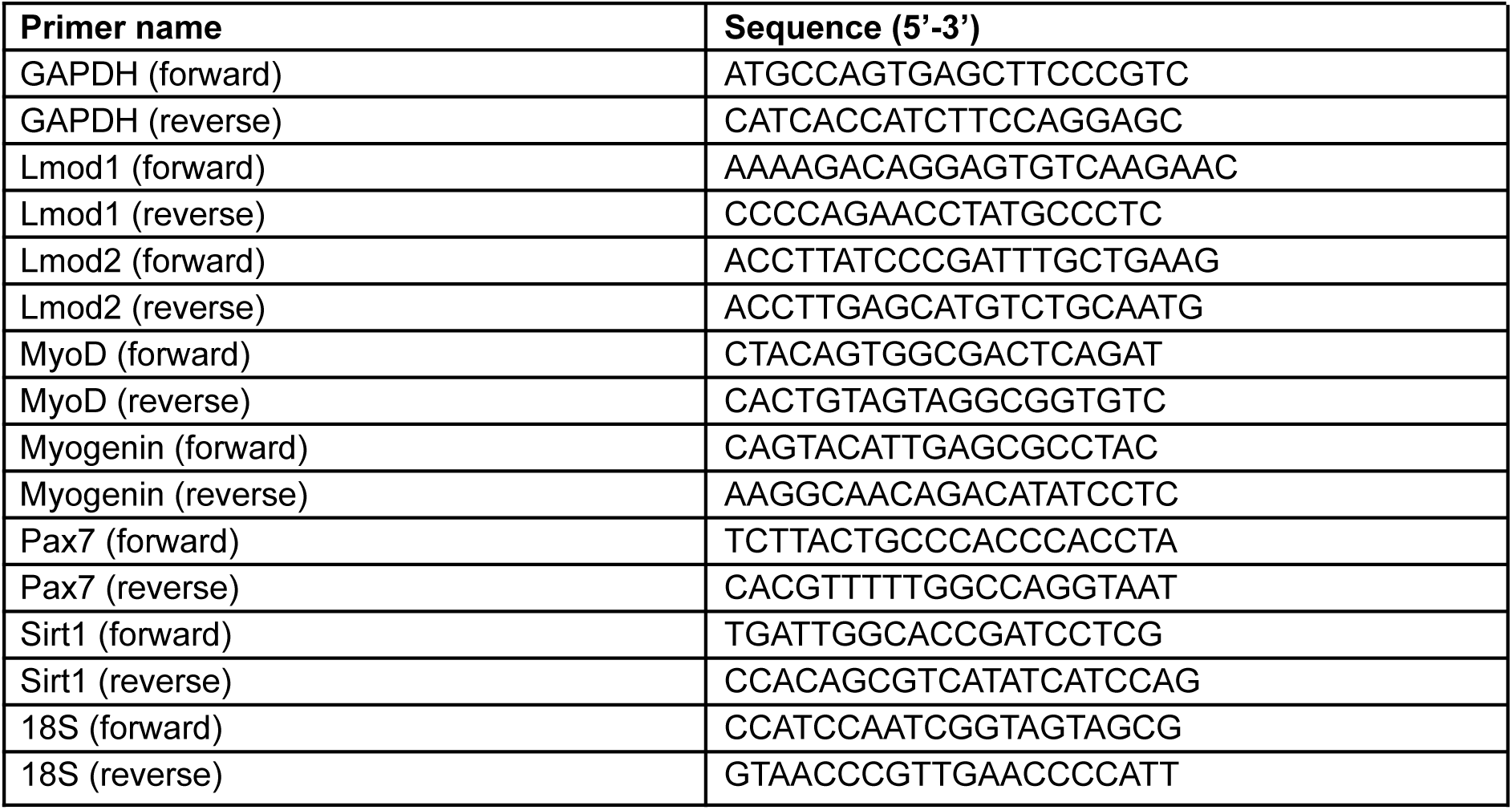
Primers for qRT-PCR.

**Table 3:** Antibody-List.

| **Antibody name** | **Source** | **Product number** |
| --- | --- | --- |
| Monoclonal rat IgG2a Sca1-FITC | eBioscience | 11-5981-85RRID:AB_465334 |
| Monoclonal rat IgG2a Sca1-PE (Ly-6A/E-PE) | BDBioscience | 553108RRID:AB_394629 |
| Hybridoma mouse IgG1 Pax7 | DSHB | Pax7RRID:AB_528428 |
| Polyclonal rabbit IgG Lmod1 | Proteintech | 15117-1-AP |
| Polyclonal human IgG Lmod2 | Atlas Antibodies | HPA051039 |
| Polyclonal chicken IgG Laminin | LSBio | LS-C96142 |
| Hybridoma mouse IgG1 myogenin | DSHB | F5D |
| Hybridoma mouse IgG1 devMHC | DSHB | MF20 |
| Polyclonal rabbit IgG Ki-67 | Abcam | ab15580 |
| Monoclonal mouse IgG2b Sirt1 | Proteintech | 60303-1-Ig |
| bisBenzimideH 33258 (Hoechst) | Sigma-Aldrich | B2261 |
| Monoclonal mouse M2 Flag | Sigma-Aldrich | F1804 |
| Monoclonal mouse IgG GFP | Santa Cruz | Sc-9996 |
| Monoclonal mouse IgG BirA | Novus Biologicals | 5B11c3-3 |
| Monoclonal mouse IgG1 Gapdh | SantaCruz | sc-365062 |
| Monoclonal mouse IgG H4 (L64C1) | Cell Signaling | #2935 |
| Monoclonal mouse IgG Aldoa | Proteintech | 67453-1-Ig |
| anti-mouse IgG Alexa Fluor® 488 | Thermo Fisher | A-21121 |
| anti-mouse IgG Alexa Fluor® 546 | Thermo Fisher | A-21123 |
| anti-mouse IgG2b Alexa Fluor® 488 | Thermo Fisher | A-21141 |
| anti-mouse IgG2b Alexa Fluor® 488 | Thermo Fisher | A-21242 |
| anti-rabbit IgG Alexa Fluor® 488 | Thermo Fisher | A-21206 |
| anti-rabbit IgG Alexa Fluor® 546 | Thermo Fisher | A10040 |
| anti-rabbit IgG Alexa Fluor® 647 | Thermo Fisher | A-31573 |
| anti-chicken IgG Alexa Fluor® 488 | Thermo Fisher | A-11039 |
| Anti-Streptavidin Alexa Fluor® 568 | Thermo Fisher | S11226 |
| Anti-Mouse Immunoglobulins/HRP | Agilent Dako | P0448 |
| Anti-Rabbit Immunoglobulins/HRP | Agilent Dako | P0447 |

### Immunofluorescence staining of HEK293T cells

The cells were washed 3x with 1 x PBS and fixed in 4 % formaldehyde (v/v) in PBS for 10 minutes at room temperature. After that, cells were washed again 3x with 1 x PBS and permeabilized using a permeabilization buffer (0.7 % Triton X-100 in 1 x PBS) for 15 minutes at room temperature, followed by two washing steps with PBS for 5 minutes each. The samples were incubated with a blocking solution (10 % (w/v) BSA, 10 % (v/v) Triton X-100, 5 % (v/v) goat serum) for 1h at room temperature. The coverslips were incubated with primary antibody anti-FLAG (Sigma-Aldrich, mouse, F1804, 1:500) at 4 °C overnight. After washing 3x with PBS, the secondary fluorescence-labeled antibody anti-mouse IgG (Alexa Fluor® 488, A-21121, Thermo Fisher, 1:1000) in blocking solution and anti-biotin (Streptavidin AlexaFuor® 568, S11226, Thermo Fisher, 1:2000) in blocking solution were incubated for 1 h at RT. After washing once with PBS nuclei were stained with Hoechst (bisBenzimideH 33258, B2261 Sigma, 0.02 μg/μl in PBS, 1:5000) at RT for 5 minutes. and were finally washed twice for 5 minutes with PBS. Finally, the samples were washed twice with PBS for 5 minutes, and cells were mounted in Permafluor mounting medium (TA-006-FM, Thermo Fisher) and coverslips (631-1574, VWR). Immunofluorescence microscopy was performed with an Axio Imager (Z2 using a Plan-Apochromat 63 x / 0.8 M27 Objective) and analyzed with Zen 2 Blue Edition software (Carl Zeiss Microscopy GmbH).

### Proximity ligation assay

Primary myoblasts were cultured in a collagen-coated removable 12-well chamber (81201, Ibid) and fixed in 2 % PFA for 5 minutes at RT. Subsequently, the proximity ligation assay (PLA) (#DUO92008, Sigma-Aldrich) was performed according to the manufacturer’s protocol. Anti-Sirt1 antibodies and anti-Lmod1 antibodies were incubated overnight at 4 °C. Immunofluorescent pictures were taken with an Axio Imager (Z2 using a Plan-Apochromat 63x / 0.8 M27 Objective) and analyzed with Zen 2 Blue Edition software (Carl Zeiss Microscopy GmbH).

### Harvesting and lysis of cells for immunoblotting

To harvest the cultured cells from proliferation or differentiation assay, the medium was removed and the cells were washed with PBS at RT. Then, 50 μl RIPA buffer (150 mM Sodium Chloride (P029.2, Roth), 1 % Triton X-100 (v/v, 3051.3, Roth), 0.5 % Sodium Deoxycholate (89904, Thermo Fisher), 0.1 % SDS (w/v, 75746-250G, Sigma), 50 mM Tris (4855.2, Roth), pH8) was added to each well of the 24 well plate. The cells were carefully scraped off the plate by using a cell scraper. Cells were incubated on ice for 20 min and mixed by vortexing every 5 min. Cell lysis was completed by sonication using Diagenode’s Bioruptor® Plus 10x for 1 min (1 min ‘on’, 30 sec ‘off’).

Primary antibodies used for immunoblotting analysis: anti-BirA (Novus Biologicals, 5B11c3-3, 1:500), anti-GFP (Santa Cruz, sc-9996, 1:1000), anti-mouse (Agilent Dako, P0448, 1:1500) anti-Lmod1 (Proteintech, 15117-1-AP, 1:500), anti-Lmod2 (Atlas Antibodies, HPA051039, 1:500), anti-Sirt1 (Proteintech, 60303-1-Ig, 1:1000), anti-Aldoa (Proteintech, 67453-1-Ig, 1:20.000), anti-H4 (Cell Signaling, #2935, 1:1000), anti-Gapdh (SantaCruz, sc-365062, 1:200). Secondary antibodies: anti-rabbit (Agilent Dako, P0447, 1:2000), anti-mouse (Agilent Dako, P0448, 1:1500).

### Protein quantification assays

The EZQ™ Protein Quantitation Kit (R33200, Thermo Fisher) and the Invitrogen™ Qubit™ 3 Fluorometer (15387293, Invitrogen), together with Qubit™ assay tubes (Q32856, Invitrogen), were used to quantify protein samples according to the manufacturer’s protocols. The TECAN microplate reader (1405005206, Tecan INFINITE M1000 PRO, excitation 485 nm, emission 580 nm) was used to analyze the EZQ assay.

### GFP Trap

For the generation of cell lines transiently expressing Sirt1 fused to GFP or GFP control, plasmids were generated using Gateway Technology (Invitrogen). Four 10 cm dishes of HEK293T cells (3 million cells/dish) were used for each candidate, and prior to transfection, the medium was replaced with a transfection medium (DMEM with 1 % FBS, without antibiotics). Each plate was transfected with 5 µg plasmid DNA and 15 µg Polyethylenimine (PEI 25K™, Polysciences, 23966-100), both previously prepared in OptiMEM (Gibco, 11520386). The transfection medium was changed to 5 mL DMEM with 10 % FBS and 1 % Pen-Strep 6 h post-transfection. Additionally, 5 µl (1 mg/mL) of Tetracycline was to the media and cells were incubated for 48 h at 3.5 % CO2, 37 °C. Subsequently, cells were harvested by trypsinization, and 20 million cells per pellet were collected for GFP-Trap pull-down. Each pellet was lysed in 200 µl of lysis buffer (20 mM Tris pH 7.5, 150 mM NaCl, 1 mM EDTA, 0.3 % Triton, 10 % Glycerol) containing protease and phosphatase inhibitors. Tubes were placed on ice for 30 min with vortexing every 10 min, followed by brief sonication in a Bioruptor Plus for 5 cycles (30 sec on/30 sec off) at the highest setting and afterwards centrifuged at 20000x *g*, for 10 min at 4 °C. Lysate-supernatants were transferred to a pre-cooled tube and to each of them 300 µl dilution buffer (10 mM Tris pH 7.5, 150 mM NaCl, 0.5 mM EDTA), was added. 25 μl of GFP Trap beads (ChromoTek GFP-Trap® Agarose, Proteintech, gta) were equilibrated in 0.5 mL of ice-cold dilution buffer and were spun down at 100x *g* for 5-10 sec at 4 °C. Beads were washed two more times with 500 μl dilution buffer. Cell lysates were added to equilibrated GFP Trap beads and incubated for 1 h at 4 °C with constant mixing. Tubes were spun at 100x *g* for 5-10 sec at 4 °C. GFP Trap beads were washed with 500 μl ice-cold dilution buffer, followed by two washes with wash buffer (10 mM Tris pH 7.5, 150 mM NaCl, 0.05 % P40 Substitute, 0.5 mM EDTA). To elute proteins, 2 x 30 µl of 2x SDS-sample buffer (120 mM Tris pH 6.8, 20 % glycerol, 4 % SDS, 0.04 % bromophenol blue) was added to the GFP Trap beads and boiled for 5 min at 95 °C. Subsequently, the beads were collected by centrifugation at 2500x *g* for 2 min and supernatant was collected to perform an SDS-PAGE.

### Nuclear cytoplasmic fractionation experiment

Nuclear cytoplasmic fractionation was performed seeding 3 x 10^6^ primary myoblasts. Prior cell harvest, cells were washed twice with 10 mL of PBS at RT to remove any residual media. Then, they were trypsinized with 0.25 % Trypsin and resuspended in 8 mL of myoblasts growth medium. After centrifuging at 500x *g* for 5 minutes, the supernatant was discarded, and the cells were resuspended in 5 mL of ice-cold PBS. At this stage, the total number of cells was counted. For hypotonic lysis, the cells were spun at 4 °C and 500x *g*. The resulting pellet was resuspended in 500-700 µl of Buffer A (10 mM Tris pH 7.5) containing protease inhibitor. The cells were incubated on ice for 20-30 minutes, and their swelling was monitored. Then, they were lysed by passing them through a 27G needle 15-30 times. The process was stopped when 70-80 % of the cells were lysed to prevent nuclei breakdown. Nuclei were manually counted to calculate the ratio of nuclei to total cells. The lysed cells were centrifuged at 1000x *g* at 4 °C to separate nuclei, and the supernatant was collected as the cytoplasmic fraction. The pellet of nuclei was resuspended in 60-100 µl of Buffer B (0.25 M sucrose, 50 mM Tris pH 7.5, 25 mM Kcl, 5 mM MgCl2, 2 mM DTT) with protease inhibitor and the nuclei were manually counted using a hemocytometer.

### RNA isolation

RNA isolation was performed with TriFast (Peqlab) according to the manufacturer’s protocol. Primary myoblasts were either directly harvested into TriFast or cell pellets were resuspended and lysed by pipetting.

### cDNA synthesis

cDNA was synthesized by reverse transcription using 600 ng of isolated RNA. The GoScript™ Reverse Transcriptase kit (A5001, Promega) was used according to the manufacturer’s protocol. Finally, the cDNA was diluted 1:10 and stored at −20 °C until further analysis.

### qRT-PCR

Quantitative real-time PCR was performed on a CFX384 Touch Real-Time PCR Detection System (Biorad). One qRT-PCR reaction mix consists of 3 µl cDNA, 5 µl iQ SYBR™ Green Supermix (1708882, Biorad), 1 µl sterile ddH2O and 0.5 µl of 10 µM forward and reverse primer (Metabion, ordered as 100 μM stock in sterile ddH2O, sequences listed in Table 2). Technical triplicates were performed for each sample. For triplicates having a Cq-SEM (calculated by Bio-Rad CFX Maestro 1.1) higher than 0.4, the replicate with the largest difference was excluded. The obtained Ct-values of target genes and housekeeping genes were used to calculate relative expression according to the 2^-ΔΔCt^ formula (Pfaffl 2001).

### RNA Sequencing

Primary myoblasts were seeded and pre-treated with siRNA against Lmod1 and Sirt1 for two days under proliferating conditions. A second transfection with siLmod1 and siSirt1 was performed simultaneously with the induction of differentiation. After one day of differentiation, the cells were collected in TriFast and RNA was isolated using the procedure described above. The concentration and integrity of the isolated RNA was determined using Qubit 3.0 Fluorometer (Thermo Fisher), Bioanalyzer 2100, and Tapestation RNA (Agilent). To prepare the poly-A library, TruSeq RNA Library Prep Kit v2 (Illumina, RS-122-2001) was used according to the manufacturer’s protocol. The process started with the enrichment of poly-A-containing mRNA using magnetic beads, followed by first-strand synthesis of cDNA using SuperScript III Reverse Transcriptase (Thermo Fisher). The second strand synthesis was performed using the second strand master mix (TruSeq RNA Library Prep Kit v2 (Illumina)), and the DNA was cleaned up using Agencourt AMPure XP Beads (Beckman Coulter, A63881). After end repair and dA-tailing, adaptors were ligated, and the library was further enriched using the PCR Master Mix (TruSeq RNA Library Prep Kit v2 (Illumina)). The concentration and quality of the library were determined using Qubit 3.0 Fluorometer (Thermo Fisher), Bioanalyzer 2100, and Tapestation DNA (Agilent). Finally, the libraries were pooled and RNA-Seq and data analysis was performed by Azenta US using Illumina NovaSeq (USA; www.GENEWIZ.com).

### Whole proteome analysis

For proteomics analysis of myoblasts or Lmod1 overexpression cell lines, samples were sonicated (Bioruptor Plus, Diagenode, Belgium) for 10 cycles (60 sec ON/30 sec OFF) at high setting, at 20 °C, followed by boiling at 95 °C for 5 min. Reduction with 10 mM Dithiothreitol (DTT) (6908.3, Roth) for 15 min was followed by alkylation with iodoacetamide (IAA, final concentration 15 mM) for 30 min at room temperature in the dark. Protein amounts were estimated following an SDS-PAGE gel of 10 µL of each sample against an in-house cell lysate of known quantity. 30 µg of each sample was used for digestion. Proteins were precipitated overnight at −20 °C after addition of 4x volume of ice-cold acetone. The following day, the samples were centrifuged at 20800x *g* for 30 min at 4 °C and the supernatant was carefully removed. Pellets were washed twice with 300 µL ice-cold 80 % (v/v) acetone in water and then centrifuged at 20800x *g* at 4 °C for 10 min. After removing the acetone, pellets were air-dried before addition of 25 µL of digestion buffer (1M Guanidine, 100 mM HEPES, pH 8). Samples were resuspended with sonication as explained above, then LysC (Wako) was added at 1:100 (w/w) enzyme:protein ratio and digestion proceeded for 4 h at 37 °C under shaking (1000 rpm for 1 h, then 650 rpm). Samples were then diluted 1:1 with MilliQ water and trypsin (Promega) added at at 1:100 (w/w) enzyme:protein ratio. Samples were further digested overnight at 37 °C under shaking (650 rpm). The day after, digests were acidified by the addition of TFA to a final concentration of 10 % (v/v), heated at 37 °C and then desalted with Waters Oasis® HLB µElution Plate 30 µm (Waters Corporation, MA, USA) under a soft vacuum following the manufacturer instruction. Briefly, the columns were conditioned with 3×100 µL solvent B (80 % (v/v) acetonitrile; 0.05 % (v/v) formic acid) and equilibrated with 3×100 µL solvent A (0.05 % (v/v) formic acid in Milli-Q water). The samples were loaded, washed 3 times with 100 µL solvent A, and then eluted into 0.2 mL PCR tubes with 50 µL solvent B. The eluates were dried down using a speed vacuum centrifuge (Eppendorf Concentrator Plus, Eppendorf AG, Germany). Dried samples were stored at −20 °C until analysis.

### Enrichment and digestion of biotinylated proteins (BioID)

The protocol was used as described in Bartolome et al (Bartolome et al. 2023). In short, pellets corresponding to 20 Mio cells were resuspended in 4.75 mL lysis buffer (50 mM Tris pH 7.5; 150 mM NaCl; 1 mM EDTA; 1mM EGTA; 1 % (v/v) Triton X-100; 1 mg/ml aprotinin; 0.5 mg/ml leupeptin; 250 U turbonuclease; 0.1 % (w/v) SDS) and incubated for 1 h at 4 °C. Streptavidin Sepharose High Performance (GE Healthcare) was acetylated by the addition of 10 mM sulfo-NHS acetate (Thermo Fisher Scientific) for 30 min twice and then equilibrated in lysis buffer. For each lysate 80 µl of equilibrated beads were used. Upon loading, beads were washed 5 times with 600 µl 50 mM AmBic. On-bead digest was performed with 200 µl LysC (5 ng/µl) at 37 °C overnight. First elution step was achieved using two times 150 µl of 50 mM AmBic. After pooling both fractions, peptides were further digested off-beads by adding 1 µg trypsin and incubating at 37 °C for 3 h. Biotinylated peptides were eluted in a second elution step using two times 150 µl 20 % TFA (Biosolve) in acetonitrile (Biosolve). Both fractions were pooled and neutralized to pH 8.0 by adding 50 µl 200 mM HEPES and sodium hydroxide as necessary. Peptides were digested further off-beads by adding 1 µg trypsin at 37 °C for 3 h. Digested peptides from the two elution steps were desalted using Waters Oasis® HLB μElution Plate 30 μm (Waters) according to the manufacturer’s instructions.

### LC-MS/MS analysis

Peptides were reconstituted in 10 µl reconstitution buffer at a concentration of 1 µg/µl, 0.5 μl of the HRM kit (Ki-3002-1, Biognosys) was added in a dilution recommended by the manufacturer and 1 μl were injected for measurement. Peptides were separated using the UltiMate 3000 UPLC system (Thermo Fisher Scientific, for myoblasts) or nanoAcquity UPLC system (Waters, for BioID and Lmod1 overexpressing cell lines) fitted with trapping (nanoAcquity Symmetry C18, 5µm, 180 µm x 20 mm) and an analytical column (nanoAcquity BEH C18, 1.7µm, 75µm x 250mm). The outlet of the analytical column was coupled directly to Q-exactive HF (Thermo Fisher Scientific, for myoblasts) or Orbitrap Exploris 480 (Thermo Fisher Scientific, for BioID and Lmod1 overexpressing cell lines) using the Proxeon nanospray source. For myoblast experiment, solvent A was water, 0.1 % FA and solvent B was 80 % (v/v) acetonitrile, 0.08 % FA. Peptides were eluted via a non-linear gradient from 1 % to 62.5 % B in 131 min. Total runtime was 150 min, including clean-up and column re-equilibration. The peptides were introduced into the mass spectrometer via a Pico-Tip Emitter 360 µm OD x 20 µm ID; 10 µm tip (New Objective) and a spray voltage of 2.2 kV was applied. The RF ion funnel was set to 60 %.

Data Dependent Acquisition (DDA) settings were as follows: Full scan MS spectra with mass range 350-1650m/z were acquired in the Orbitrap with a resolution of 60,000 FWHM. The filling time was set at a maximum of 20ms with an AGC target of 3×10^6^ ions. A Top15 method was employed to select precursor ions from the full scan MS for fragmentation (minimum AGC target of 1x 10^3^ ions, normalized collision energy of 27 %), quadrupole isolation (1.6 m/z) and measurement in the Orbitrap (resolution 15,000 FWHM, fixed first mass 120 m/z). The fragmentation was performed after the accumulation of 2×10^5^ ions or after filling time of 25ms for each precursor ion (whichever occurred first). Only multiply charged (2+ −7+) precursor ions were selected for MS/MS. Dynamic exclusion was employed with a maximum retention period of 20s. Isotopes were excluded.

For Data Independent Acquisition (DIA), 1 μg of reconstituted peptides were loaded using the same setup and LC conditions used for DDA. MS conditions were modified as follows: Full scan MS spectra with mass range 350-1650 m/z were acquired in profile mode in the Orbitrap with resolution of 120,000 FWHM. The filling time was set at a maximum of 60 ms with an AGC target of 3×10^6^ ions. DIA scans were acquired with 40 mass window segments of differing widths across the MS1 mass range. The default charge state was set to 3+. HCD fragmentation (stepped normalized collision energy; 25.5, 27, 30 %) was applied and MS/MS spectra were acquired with a resolution of 30,000 FWHM with a fixed first mass of 200 m/z after accumulation of 3x 10^6^ ions or after a filling time of 35ms (whichever occurred first). Data was acquired in profile mode. For data acquisition and processing Tune version 2.9 Q-Exactive HF For BioID project experiment and, solvent A was water, 0.1 % FA and solvent B was 80 % (v/v) acetonitrile, 0.08 % FA. Peptides were eluted via a non-linear gradient from 3 % to 40 % B in 90 min. Total runtime was 115 min, including clean-up and column re-equilibration. The peptides were introduced into the mass spectrometer via a Pico-Tip Emitter 360 µm OD x 20 µm ID; 10 µm tip (New Objective) and a spray voltage of 2.2 kV was applied. The RF ion funnel was set to 30 %. For Data Independent Acquisition (DIA), MS conditions were modified as follows: Full scan MS spectra with mass range 350-1650 m/z were acquired in profile mode in the Orbitrap with resolution of 120,000 FWHM. The filling time was set at a maximum of 60 ms with an AGC target of 3×10^6^ ions (300 %). DIA scans were acquired with 34 mass window segments of differing widths across the MS1 mass range. The default charge state was set to 2+. HCD fragmentation (stepped normalized collision energy; 25.5, 27, 30 %) was applied and MS/MS spectra were acquired with a resolution of 30,000 FWHM with a fixed first mass of 200 m/z after accumulation of 3x 10^6^ ions (3000 %) or after a filling time of 40ms (whichever occurred first). Data was acquired in profile mode. For data acquisition and processing Tune 3.1.

For Lmod1 overexpressing cell line experiment, solvent A was water, 0.1 % FA and solvent B was acetonitrile, 0.1 % FA. Peptides were eluted via a non-linear gradient from 3 % to 40 % B in 120 min. Total runtime was 145 min, including clean-up and column re-equilibration. The peptides were introduced into the mass spectrometer via a Pico-Tip Emitter 360 µm OD x 20 µm ID; 10 µm tip (New Objective) and a spray voltage of 2.2 kV was applied. The RF ion funnel was set to 30 %. For Data Independent Acquisition (DIA), 1 μg of reconstituted peptides were loaded using the same setup and LC conditions used for DDA. MS conditions were modified as follows: Full scan MS spectra with mass range 350-1650 m/z were acquired in profile mode in the Orbitrap with resolution of 120,000 FWHM. The filling time was set at a maximum of 60 ms with an AGC target of 3×10^6^ ions. DIA scans were acquired with 40 mass window segments of differing widths across the MS1 mass range. The default charge state was set to 3+. HCD fragmentation (stepped normalized collision energy; 25.5, 27, 30 %) was applied and MS/MS spectra were acquired with a resolution of 30,000 FWHM with a fixed first mass of 200 m/z after accumulation of 3x 10^6^ ions or after a filling time of 35 ms (whichever occurred first). Data was acquired in profile mode. For data acquisition and processing Tune 2.0.

### Proteomics data analysis

For myoblasts DIA search against a library was performed. Acquired data were processed using Spectronaut Professional v13 (Biognosys AG). For library creation, the DDA and DIA raw files were searched with Pulsar (Biognosys AG) against the mouse UniProt database (*Mus musculus,* v. 160106, 16,747 entries) with a list of common contaminants (247 entries) Swissprot database appended, using default settings. For library generation, default BGS factory settings were used.

DIA data were then uploaded and searched against this spectral library using Spectronaut Professional (v.10 and v.13 Biognosys) and default settings. Relative quantification was performed in Spectronaut for each pairwise comparison using a two-sided t-test performed at the precursor level followed by multiple testing correction and default settings, except: Major Group Quantity = median peptide quantity; Major Group Top N = OFF; Minor Group Quantity = median precursor quantity; Minor Group Top N = OFF; Data Filtering = Q value sparse; Normalization Strategy = Local normalization; Row Selection = Q value complete.

For Lmod1 overexpression cell lines, DIA raw data were analyzed using the directDIA pipeline in Spectronaut v.14 (Biognosysis, Switzerland) with BGS settings besides the following parameters: Protein LFQ method= QUANT 2.0, Proteotypicity Filter = Only protein group specific, Major Group Quantity = Median peptide quantity, Minor Group Quantity = Median precursor quantity, Data Filtering = Qvalue, Normalizing strategy = Local Normalization. The data were searched against an in-house database (*Mus Musculus*, 16,747 entries) and a contaminants (247 entries) Swissprot database. The data were searched with the following variable modifications: Oxidation (M) and Acetyl (Protein N-term). A maximum of 2 missed cleavages for trypsin and 5 variable modifications were allowed. The identifications were filtered to satisfy FDR of 1 % on peptide and protein level. Relative quantification was performed in Spectronaut for each paired comparison using the replicate samples from each condition. The data (candidate table) and data reports (protein quantities) were then exported, and further data analyses and visualization were performed with Rstudio using in-house pipelines and scripts.

BioID DIA raw data were analyzed using the directDIA pipeline in Spectronaut with v.15. The data were searched against a specific species (*Homo sapiens,* v. 160112, 20,375 entries) and a contaminants (247 entries) Swissprot database. The data were searched with the following modifications: Oxidation (M), Acetyl (Protein N-term) and Biotin_K. A maximum of 2 missed cleavages for trypsin and 5 variable modifications were allowed. The identifications were filtered to satisfy FDR of 1 % on peptide and protein level. Relative quantification was performed in Spectronaut using LFQ QUANT 2.0 method with Global Normalization, precursor filtering percentile using fraction 0.2 and global imputation. The data (candidate table) and data reports (protein quantities) were then exported and further data analyses and visualization were performed with Rstudio using in-house pipelines and scripts.

### Clustering and gene set enrichment analysis

Proteomics data for the myoblast differentiation time course were filtered for protein groups quantified by at least two unique (proteotypic) peptides. All the differentiation time points (1d:5d) were compared to undifferentiated myoblast (0d) and protein groups that showed an absolute AVG Log2 Ratio > 0.58 and Q value < 0.25 at any time point were selected for clustering analysis. Determination of an optimal number of clusters, clustering, and enrichment analysis for each cluster was performed with ClueR (Yang et al. 2015) using the AVG Log2 Ratios to 0d as input.

A gene set was created to analyze the LMOD1 overexpression cell line using all the protein groups differentially abundant in 1d vs. 0d myoblast differentiation time course (AVG Log2 Ratio > 0.58 and Q value < 0.05). This gene set was used as input for a gene set enrichment analysis based on AVG Log2 Ratios from the comparison of LMOD1 OE vs. GFP OE performed with WebGestalt (Liao et al. 2019).

### Statistical analysis

Mouse and cell line experiments were performed at least in biological triplicates, and numbers are indicated in the figure legends. The results are shown as the mean ± SEM unless indicated otherwise in the figure legends. Statistical significance was calculated using the GraphPad Prism software using the statistical test indicated in the figure legends, with *: p 0.05, **: p 0.01, ***: p 0.001, ****: p 0.0001 and ns (not significant) or nd (not detectable). RNA-seq approaches were conducted in biological replicates (n = 5) using primary myoblasts isolated from individual mice per experimental group, and statistical analysis is described in the respective section.

## Data availability

The mass spectrometry proteomics data have been deposited to the MassIVE (https://massive.ucsd.edu) repository with the following dataset identifier:

Proteomic dataset: MSV000093379
Username: MSV000093379_reviewer
Password: ES_Lmod1

RNA-Seq data have been deposited on Gene Expression Omnibus (GEO) international public repository at: GSE254443
Password: epkbcyyinjmxzmd

## Notes

### Competing Interest Statement

The authors have declared no competing interest.

### Summary of Updates

Strengthened in vivo validation of the LMOD1-SIRT1 axis. We added new immunostainings for SIRT1 and PAX7 to our muscle regeneration time course. Analysis at day 5 post-injury, coinciding with peak LMOD1 expression, demonstrates increased SIRT1 abundance and localization in mononucleated cells and newly formed myofibers (revised Figure 4I), supporting the in vivo relevance of our proposed mechanism. Enhanced rigor and antibody validation. Antibody specificity for LMOD1 and SIRT1 was validated using siRNA knockdown and overexpression approaches, including new immunofluorescence controls. We clarified that proximity ligation assays demonstrating the LMOD1-SIRT1 interaction were performed in primary myoblasts. The discussion was refined to better articulate a dynamic, stage-dependent model of LMOD1 function during differentiation. Clarified cellular phenotypes following Lmod1 knockdown. To address concerns regarding cell health, we performed new TUNEL assays showing no increase in apoptosis upon Lmod1 depletion (new Figure S2I). These data support the conclusion that LMOD1 loss impairs differentiation rather than inducing cell death. Variability in morphology was addressed by emphasizing the use of multiple independent primary myoblast cultures and additional quantitative marker analyses. Improved proteomic and comparative analyses. We added quantification of LMOD1- and LMOD2-BirA bait proteins to validate BioID comparability and highlighted SIRT1 in revised volcano plots. We also incorporated analyses of published transcriptomic datasets to clarify the temporal regulation of Lmod1 during muscle stem cells activation to strengthen the rationale for focusing on LMOD1 relative to other leiomodin isoforms.

